# High resolution photocatalytic mapping of SARS-CoV-2 Spike protein-host cell membrane interactions

**DOI:** 10.1101/2022.09.02.506438

**Authors:** Suprama Datta, Alexander H. Tavares, Tamara Reyes-Robles, Keun Ah Ryu, Nazimuddin Khan, Tyler J. Bechtel, Jayde M. Bertoch, Cory H. White, Daria J. Hazuda, Kalpit A. Vora, Erik C. Hett, Olugbeminiyi O. Fadeyi, Rob C. Oslund, Mohsan Saeed, Andrew Emili

## Abstract

Identifying protein environments at the virus-host cell interface can improve our understanding of viral entry and pathogenesis. SARS-CoV-2, the virus behind the ongoing COVID-19 pandemic, uses the cell surface ACE2 protein as a major receptor, but the contribution of other cellular proteins in the entry process is unknown. To probe the microenvironment of SARS-CoV-2 Spike-ACE2 protein interactomes on human cells, we developed a photocatalyst-based viral-host protein microenvironment mapping platform (ViraMap) employing iridium photocatalysts conjugated to Spike for visible-light driven proximity labelling on host cells. Application of ViraMap on ACE2-expressing cells captured ACE2, the established co-receptor NRP1, as well as other proteins implicated in host cell entry and immunomodulation. We further investigated these enriched proteins via loss-of-function and over-expression in pseudotype and authentic infection models and observed that the Ig receptor PTGFRN and tyrosine kinase ligand EFNB1 can serve as SARS-CoV-2 entry factors. Our results highlight additional host targets that participate infection and showcase ViraMap for interrogating virus-host cell surface interactomes.

## Introduction

The pathogenesis and life cycle of the enveloped novel beta-coronavirus SARS-CoV-2 begins when its Spike protein attaches to several receptors on a host cell membrane. While these trimeric spike-like glycoprotein moieties protruding from the viral envelop serve as the viral attachment ligand, the cell membrane poses a critical biological barrier to the virus access into the host cell cytosol. Therefore, it is crucial to understand the interactions between the Spike protein and the membrane-tethered host factors at the first point of contact. However, the spectrum of membrane-associated proteins that are targeted for cell entry are not well known and may be contextual and transient in nature. While high-affinity binding is typically essential for cell entry, other lower-affinity interactions are thought to impact tissue tropism, pathogen recognition and immune signaling, adding complexity to the molecular landscape of the virus-host cell interface. Virus “co-receptors” and “activators” include a dynamic array of biochemically diverse membrane-tethered host factors broadly categorized as immunoglobulin superfamily (IgSF) members, scavenger receptors, glycans, cellular adhesion molecules (CAMs) and the phosphatidylserine (PtdSer) family of tyrosine kinase receptors (Dermody et al., 2009; Hantak et al., 2019; Maginnis, 2018; Taban et al., 2022). Some of the known host factors are distributed in highly organized cell membrane microdomains that orchestrate viral entry and exit (Hantak *et al*., 2019; Ripa et al., 2021), but the biophysical associations of others are not as clear. Furthermore, given their accessibility to systemically delivered drugs, membrane-tethered virus co-receptors represent particularly attractive candidates for anti-viral therapeutics.

In the case of SARS-CoV-2, the initial association with the host is thought to begin as low-affinity, transient interactions with ubiquitous components of the host cell membranes followed by high-affinity binding events that lead to irreversible conformational changes in the viral attachment proteins (Shilts et al., 2021), most notably upon engagement with the host angiotensin converting enzyme (ACE2) receptor that is preferentially expressed in lung tissue (Hoffmann et al., 2020; Stehle and Dermody, 2004; Zhou et al., 2020). These trimeric spike protein assemblies undergo profound conformational changes after high-affinity binding to ACE2 (Hoffmann *et al.*, 2020; Walls et al., 2020). While binding to ACE2 has been extensively characterized, less is known about the possible role of other host factors that comprise the transient interactome of the SARS-CoV-2 Spike protein on the host-cell surface. Identification of these putative co-factors can potentially explain the susceptibility of cells with low ACE2 levels (Chen et al., 2021; Li et al., 2020) and illuminate pattern-recognition associated mechanisms driving virus cell entry.

Established experimental studies aimed at the identification of SARS-CoV-2-host protein interactions have generally relied on immunoprecipitation (IP) or affinity tagging/purification (AP) combined with liquid chromatography-tandem mass spectrometry (LC-MS/MS)-based detection. These approaches preferentially detect the most stable and abundant host binding partners, while lower-affinity membrane-bound partners that are difficult to solubilize in a native conformation are potentially lost. Indeed, in one of the largest SARS-CoV-2-host interactome AP/MS-based studies which identified 332 high-confidence host interactions for 26 out of 29 SARS-CoV-2 viral proteins (Gordon et al., 2020), only 10% of the host proteins putatively bound by SARS-CoV-2 were membrane-tethered. Notably, only palmitoyl-transferase, which catalyzes the addition of palmitate onto various membrane-associated protein substrates, was captured as an interactor with SARS-CoV-2 Spike protein. Therefore, it is crucial to devise more sensitive and effective methods to map connections between the Spike and membrane-tethered host factors.

One promising approach is proximity-based protein labelling, which involves fusion of a modifying enzyme (e.g. BioID, APEX) to a target protein of interest, allowing for the covalent biotinylation on tyrosine or lysine residues of neighboring proteins (Qin et al., 2021). These methods are capable of promiscuously capturing protein environments within cellular compartments and typically rely on the genetic manipulation of the target of interest to express the cell-tagging enzyme. However, the physical alteration of the target protein with the labeling enzyme can have downstream consequences that effect protein expression and function. Indeed, BioID was recently used to probe SARS-CoV-2 viral protein proximity interactions through genetic fusion to BioID but the Spike-BioID protein fusion variant could not be stably expressed (May et al., 2022). The enzyme labeling approaches have also recently been applied within the replication and transcription complex (RTC) of murine coronavirus (MHV) where APEX2/BioID was used to map host proteins associated with RTC. The result of which led to the identification of over 500 candidate cellular proximal protein partners (V’Kovski et al., 2019), complicating downstream study of viral-related functions. Collectively, these observations highlight the need for a viral-host proximity labeling platform that offers high spatiotemporal resolution.

Visible light-mediated photocatalysis has emerged as an attractive approach for protein proximity labeling with high resolution (Ryu et al., 2021). We recently disclosed an antibody-iridium photocatalyst (IrPC) conjugate method for targeted, visible light activation of a biotin-labeled diazirine probe by means of short-range Dexter energy transfer (Geri et al., 2020). This generates a highly reactive carbene species that, compared to enzyme labeling methods, covalently labels biomolecules in a residue agnostic fashion through direct insertion into C-H, O-H, or N-H bonds. Due to fast quenching and the short half-life of the carbene intermediate (T_1/2_ < 1 ns) the activated probe is unable to diffuse farther than 4 nm (Geri *et al*., 2020). This drives a shorter labeling radius and greater spatiotemporal control compared to enzyme methods with higher resolution protein microenvironment snapshots (Oakley et al., 2022). Furthermore, the bioconjugation of a small molecular weight photocatalyst (< 1000 Da) to the targeting modality of interest results in a smaller perturbation compared to attachment of a larger enzyme.

In this study, we disclose a proteomic-based **vira**l-host protein microenvironment **map**ping platform that we refer to as **ViraMap** that relies on conjugation of a photocatalyst to the spike protein trimer of SARS-CoV-2 for targeted tagging and identification of spike-host protein interactions (**Figure 1a**). Using this approach, we identified several novel spike co-receptors and investigated their functional significance by performing both CRISPR-mediated loss-of-function and lentiviral overexpression in SARS-CoV-2 permissive human cell lines to observe their impact on virus entry in both pseudotype and true viral infection. Our successful implementation and functional validation of the ViraMap technology suggests this platform could be used to map other virus-host cell surface interaction interfaces in a high-resolution manner, irrespective of binding affinities or interactome complexities.

**Figure 1.**
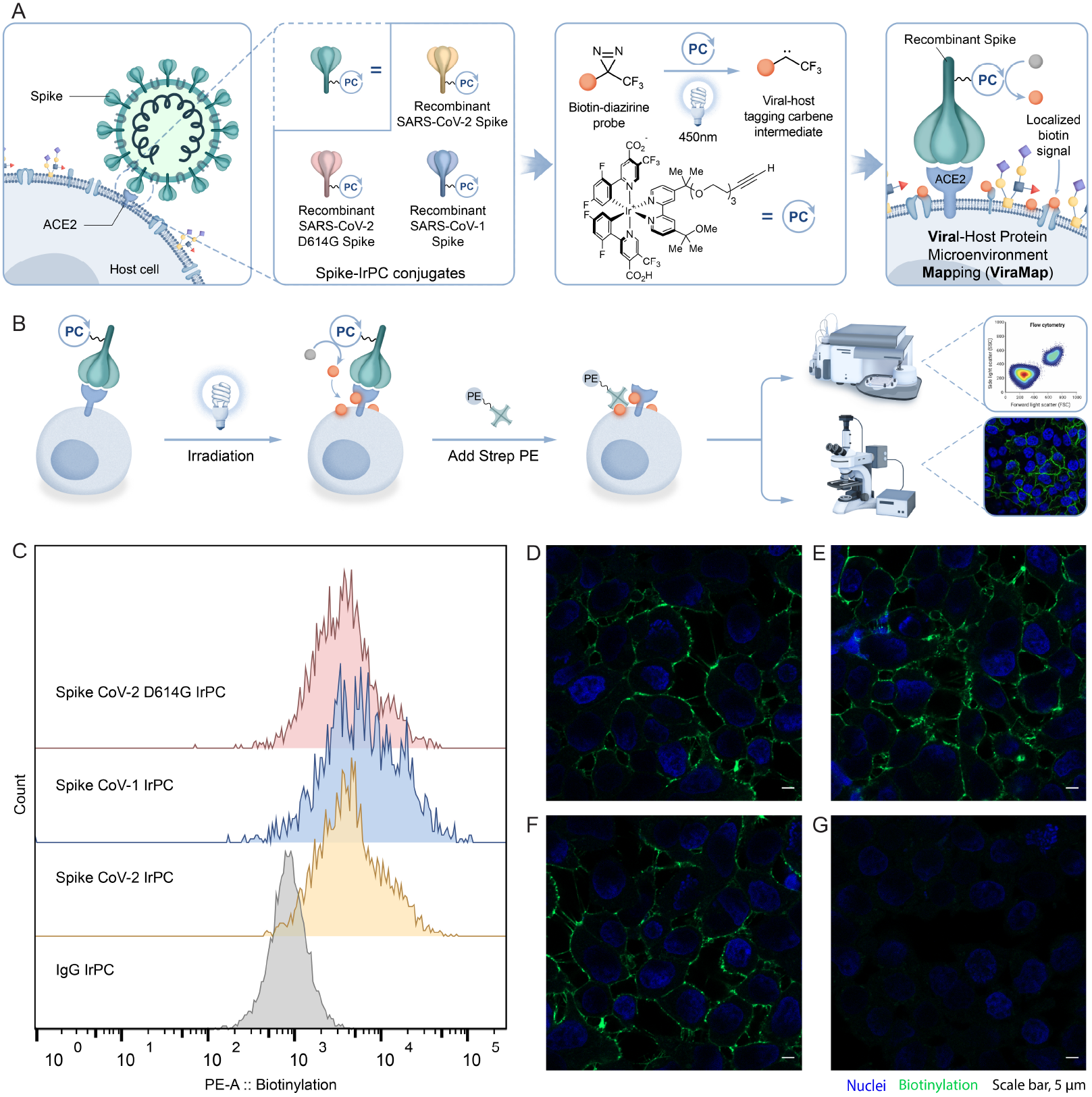
A.ViraMap study design: Selective labeling of host factors after exposing IrPC-conjugated SARS-CoV-2 Spike protein trimers, and related variants (SARS-Cov-1 Spike and SARS-CoV-2 D614G Spike) to cells expressing hACE2 under visible (450 nm) light in the presence of a biotin-diazirine probe. B. Schematics of IrPC induced labeling workflow and data analysis by flow cytometry and confocal imaging. C. Flow cytometry analysis showing cell surface biotinylation caused by the Spike-IrPC conjugate on HEK293+ACE2 cells. D. Confocal images probing the biotinylated HEK293+ACE2 cell surface upon treatment with IrPC conjugated SARS-CoV-2 D614G Spike, E. SARS-CoV-1 Spike, F. SARS-CoV-2 Spike and G. negative control isotype IgG.

## Results

### Spike-photocatalyst conjugate labels hACE2 receptor expressed on the host cell surface upon visible light irradiation

To profile SARS-CoV-2 spike-host cell surface microenvironment interactions using ViraMap, we prepared spike protein photocatalyst conjugates for targeted labeling on ACE2 expressing cells. For this we selected the SARS-CoV-2 Spike protein trimer and its related variants (SARS-CoV-1 Spike and SARS-CoV-2 D614G Spike) for conjugation with IrPC (**Figure 1a, S1a**). This was done by treating purified recombinant Spike protein variants with azidobutyric acid N-hydroxysuccinimide ester to label lysine residues with an azide group, followed by copper-catalyzed azide-alkyne click-chemical conjugation to an alkyne-bearing IrPC (**Figure S1b**). Through this bioconjugation approach, we obtained a stoichiometric ratio of the resulting Spike:IrPC conjugate of ~1:5 (**Figure S1b, Methods section**). For targeted labeling experiments, we chose HEK293T cells over-expressing the ACE2 protein (HEK293T+ACE2) as the prototype for these microenvironment profiling tests to leverage the high spike-ACE2 binding due to the elevated expression of ACE2 on the cell surface as confirmed by flow cytometry (**Figure S1c**).

With this ACE2 expressing cell line and photocatalyst-labeled Spike protein conjugates in hand, we next performed targeted labeling by exposing the cells to Spike-IrPC conjugates in the presence of a biotin-diazirine probe followed by irradiation with blue light and subsequent monitoring by flow cytometry and confocal imaging analysis (**Figure 1b**). Accordingly, HEK293T+ACE2 cells were incubated with Spike-IrPC at 4 °C to ensure selective labeling of host cell surface interactions and avoid cellular internalization of the Spike-IrPC conjugates. Flow cytometry analysis showed a clear shift in the biotinylation signal for cells treated with Spike-IrPC variant conjugates when compared with a non-binding antibody IgG control conjugated with IrPC (mouse-IgG-IrPC conjugate) (**Figure 1c**). To confirm that the Spike-IrPC conjugate retained its affinity for ACE2, we measured cell surface binding in the presence of excess free ACE2 protein (ACE2-Fc) and observed that the Spike-IrPC conjugate could be competed away from the cell surface (**Figure S1d**). Confocal imaging of Spike-IrPC induced labeling showed localization of biotin to the host cell surface environment compared to the mouse-IgG-IrPC control (**Figures 1d-g**). In addition, given that the recombinant Spike construct contains a poly-His-tag, targeted labeling with an anti-His antibody IrPC conjugate could also be used to achieve cellular biotinylation. (**Figure S1e** and **S2**).

### ViraMap elucidates low-affinity host cell membrane factors that regulate viral entry and antiviral immunity

Structurally similar spike variants are expected to interact with same set of host proteins on the cell surface. We therefore rationalized that the overlapping hits from different Spike-IrPC variants are more likely to represent bona-fide spike-interaction partners. Thus, we utilized three Spike-IrPC variants (SARS-CoV-1 spike, SARS-CoV-2 spike, and SARS-CoV-2 spike with D614G substitution) for cell surface proximity labeling on HEK293T+ACE2 cells for downstream proteomic analysis (**Figure 2a**). Furthermore, to help discern between host proteins that interact exclusively with the Spike protein versus those that are proximal to ACE2, we also performed targeted labeling using a primary-secondary antibody system consisting of an anti-ACE2 primary antibody and a secondary antibody-photocatalyst (IrPC) conjugate. The biotinylated host proteins from these targeted labeling experiments were quantitatively identified using tandem mass tag (TMT)–based LC-MS/MS quantification after isolation from the cell membrane fraction followed by streptavidin enrichment (**Figure 2a**). We confirmed the enrichment of the biotinylated proteins by western blot (**Figure S2a**).

**Figure 2.**
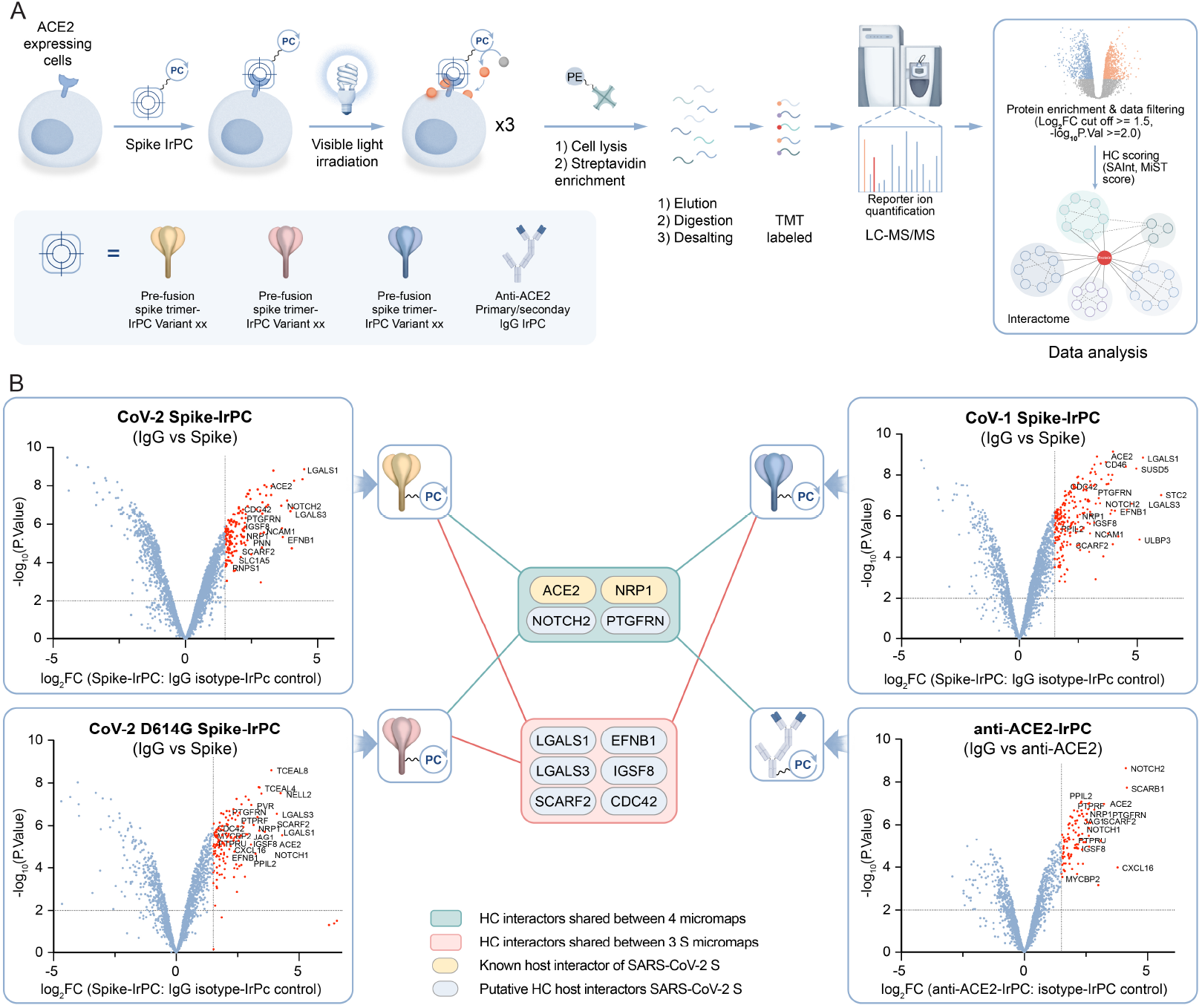
A. IrPC-labeled SARS CoV-2 Spike directed proteomics workflow. B. Enriched host factors (highlighted in red, log_2_FC≥1.5, -Log_10_Pvalue>=2.0) from targeted labeling of SARS Cov-2 wildtype Spike, SARS-CoV-2 D614G Spike, SARS-CoV-1 Spike and anti-ACE2 represented as volcano plots. The network represents an overlap of an enriched subset of proteins across proteomics experiments of 3 variants of Spike and ACE2 directed targeted labeling after scoring protein-protein interactions (MiST score cut off ≥ 0.8, SAInt FDR cut off ≤ 0.3). The nodes represent the proteins and edges represent the average of MiST score and BFDR for each interaction in the network. The nodes were clustered in 2 groups (red and blue nodes) based on the interaction overlap in the Spike-ACE2 environment. Identified host proximal proteins are denoted as yellow (known Spike proximal interactors) and blue (putative Spike proximal interactors) sub-nodes.

Targeted host cell labeling from IrPC-conjugated SARS-CoV-1 Spike, SARS-CoV-2 Spike, SARS-CoV-2 D614G Spike, and anti-ACE2 primary/secondary antibody system resulted in statistically significant enrichment of discrete sets of cellular proteins relative to the IrPC-isotype conjugate as a negative control (**Figure 2b** & **S2b**). We identified 96 high-confidence enriched proteins across all Spike-IrPC labeling experiments by scoring the protein-protein interactions using SAINTexpress (significance analysis of interactome) (Teo et al., 2014) and MiST (mass spectrometry interaction statistics) scoring algorithms (Jager et al., 2011; Verschueren et al., 2015) and applying a stringent filter (logFC ≥1.5, -log_10_ P.value ≥ 2.0) (**Figure S2c**, **Table S1 for details**). Twenty-three of these 96 high-confidence enriched proteins were shared between the all three spike variants, and 2, 25 and 46 enriched proteins were unique to each of the SARS-CoV-2, SARS-CoV-2 D614G and SARS-CoV-1 Spike-IrPC spikes, respectively (**Figure S2c**). The pathway analysis of the 23 enriched proteins (**Figure S2c**) by gene ontology enrichment analysis revealed significant association with viral entry followed by immune cell differentiation and co-stimulation processes (**Figure S2d**, **Table S2**). Moreover, the tissue expression profiles of ~70% (16 out of 23) of the enriched proteins showed broad abundance in most human tissues such as lung, gastrointestinal tract, kidney, cardiac muscle, adipose, soft tissues, reproductive tissues, and brain (Uhlen et al., 2015) (**Figure S2e**), possibly implicating their potential involvement in extra-pulmonary complications of COVID-19 (Zheng et al., 2021). Ten of the 23 high-confidence enriched proteins could be further classified into two groups based on whether they were also co-enriched with the anti-ACE2 primary/secondary antibody system (**Figure 2b, S2c**).

The first group of cell surface receptors was comprised of ACE2, NRP1, PTGFRN and NOTCH-2 (**Figure 2b**). The recurrent capture of two previously identified receptors of SARS-CoV-2 Spike, namely NRP-1 and ACE2, established the accuracy and robustness in our ViraMap proteomic platform. PTGFRN (also known as EWI-F) is a member of a novel subfamily of Ig cell surface receptors. Evidence of both protein and mRNA-level expression of PTGFRN in a vast array of tissue cells, particularly in cardiomyocytes and lung fibroblasts (Uhlen *et al*., 2015), likely suggests a high susceptibility of these cell types to SARS-CoV-2 through multiple ports of entry. Finally, the presence of Notch-signaling regulator Notch-2 in all targeted labeling experiments, as well as Notch-3 in the D614G Spike variant only, suggests a mechanism for modulation of this core host pathway upon exposure to SARS-CoV-2 as these receptors are reported to play a role in both innate and T-cell mediated immune response (Vanderbeck and Maillard, 2021).

The second group of host proteins appearing in all three Spike-IrPC labeling experiments comprised of galectins (LGALS1 and LGALS3), as well as IGSF8, EFNB1, SCARF2 and CDC42 (**Figure 2b**). These proteins were uniquely identified with all spike variants but not with the primary/secondary anti-ACE2 system suggesting their association with specific binding domains of the spike variants. The reproducible identification of galectins in these datasets aligns with the findings of recent reports on lectin-facilitated infection by SARS-CoV-2 Spike leveraging a lectin binding motif on the N-terminal domain of the Spike protein (Lempp et al., 2021). Among other identified proteins such as IGSF8 (also known as EWI-2) is member of the same Ig cell surface receptor family as PTGFRN from group 1. EFNB1 acts as a membrane-bound ligand of the Eph class of tyrosine kinase receptors that play vital roles in T-cell mediated viral immunity (Darling and Lamb, 2019), and SCARF2 belongs to the scavenger receptor family which is reported to coordinate with Toll-like receptors in recognition of hepatitis C virus (Beauvillain et al., 2010; Lyu et al., 2015; Taban *et al.*, 2022). CDC42, a Rho-GTPase that regulates the cellular actin cytoskeleton, is reported to play a key role in HIV-1 entry and B-cell differentiation required for antiviral humoral immunity (Burbage et al., 2015; Swaine and Dittmar, 2015).

Additionally, 13 other high-confidence enriched proteins shared between one or more Spike variants in our dataset were also previously implicated in playing key roles in viral entry and host antiviral immunity. Enriched proteins shared by the SARS-CoV-2 spike and SARS-CoV-1 spike targeted labeling experiments consisted of NCAM1 and SLC1A5, which are members of the Immunoglobulin-like superfamily of CAMs (IgCAMs) and known to mediate viral entry among other immunomodulatory functions (**Figure S2c, Table S2**) (Scalise et al., 2018; Srivastava et al., 2020). Furthermore, the enriched proteins identified via SARS-CoV-2 D614G Spike and SARS-CoV-1 Spike targeted labeling, include DSG2, NECTIN-2, MARCKS, MARCKSL1, ARHGAP23, SLC31A1, EPHA2, MOXD1, CDC42EP1 and EFNB3 (**Figure S2c**). DSG2 is a known receptor for human adenovirus entry (Wang et al., 2011), while NECTIN-2 is a member of the IgCAMs and is reported to serve as an entry factor for poliovirus, certain mutant strains of herpes simplex virus (HSV) and pseudorabies virus (Martinez and Spear, 2001; Xu and Jin, 2010). Both MARCKS and MARCKS-related protein (MARCKSL1) regulate protein kinase C (PKC) mediated signal transduction (Lim et al., 2015) and promote migration of inflammatory cells and the secretion of cytokines (El Amri et al., 2018). ARHGAP23 is a regulator of Rho GTPase that helps in synaptic development and plays role in innate immune response (Martin-Vilchez et al., 2017). SLC31A1, a regulator of copper homeostasis, is reported to affect Influenza A/WSN/33 (H1N1) virus replication in cultured human lung A549 cells (Rupp et al., 2017). EPHA2 serves as an epithelial cell receptor for Epstein-Barr virus entry (Zhang et al., 2018). MOXD1, a copper binding protein, is deemed crucial for Hepatitis C virus (HCV) infection (Poortahmasebi et al., 2016). CDC42 effector protein 1 (CDC42EP1), a Rho-GTPase binding protein, is known to promote cellular migration. EFNB3 acts as cellular receptors for Nipah and Hendra virus attachment glycoproteins (Bowden et al., 2008). AGTRAP, a biomarker correlated with immune infiltration, was detected only in the SARS-CoV-2 Spike and SARS-CoV-2 D614G Spike targeted labeling experiments (Liu et al., 2021) (**Figure S2c**).

Although not significantly enriched in our datasets, two other previously documented interaction partners of SARS-CoV-2 Spike, namely ITGB1 and Vimentin (VIM), were detected across all modalities in agreement with previous IP-MS and immunostaining reports (Amraei et al., 2022; Suprewicz et al., 2021; Zhang et al., 2022). However, VIM has also been reported as a common contaminant by LC-MS/MS (Mellacheruvu et al., 2013).

### Functional screen of select hits from viramapping identifies PTGFRN and EFNB1 as SARS-CoV-2 entry factors

Findings from our targeted labeling experiments highlight candidate cell surface co-receptors potentially involved in viral entry and antiviral immunity. Hence, we performed functional assays to determine which of the identified host proteins participate in SARS-CoV-2 entry into cells. We focused our investigation on the 10 overlapping proteins shared by all three Spike-IrPC variants for functional validation. Since our labelling experiments utilized HEK293T+ACE2 cells, we employed these cells for the initial functional screen, followed by Caco-2 cells that endogenously express ACE2 and have been extensively used in the filed for SARS-CoV-2 studies (Bojkova et al., 2020; Pommerenke et al., 2021; Yeung et al., 2021).

We performed CRISPR-Cas9 mediated depletion of proteins in HEK293T+ACE2 cells, infected these cells with SARS-CoV-2 pseudoparticles containing the GFP and firefly luciferase reporter genes, and monitored the pseudoparticle entry by measuring the luciferase activity (**Figure 3a**). Since HEK293T+ACE2 cells overexpress ACE2, the CRISPR/Cas9-mediated knockdown (KD) only caused ~30% reduction in ACE2 abundance, precluding the use of ACE2 KD as a positive control in these experiments. (**Figure 3b** & **S3a**). Consistent with recent reports (Lempp *et al*., 2021; Zhang et al., 2021), we saw no significant change in pseudoparticle entry following NRP1 KD despite 40% reduction in protein expression compared to controls (**Figure S3a**). Knockdown of IGSF8, SCARF2 and Notch-2 also did not result in noticeable change in infectivity (**Table S3**), possibly due to abundant ACE2 availability and/or redundancy with other Notch receptors. In striking contrast, we observed significant inhibition of pseudoparticle entry in PTGFRN KD cells (75.5% inhibition, *p*-value = 0.0082) as compared to non-targeting sgRNA control (**Figure 3b**). Knockdown of EFNB1 also resulted in ~50% inhibition of entry as compared to the negative control (**Figure 3b**).

**Figure 3.**
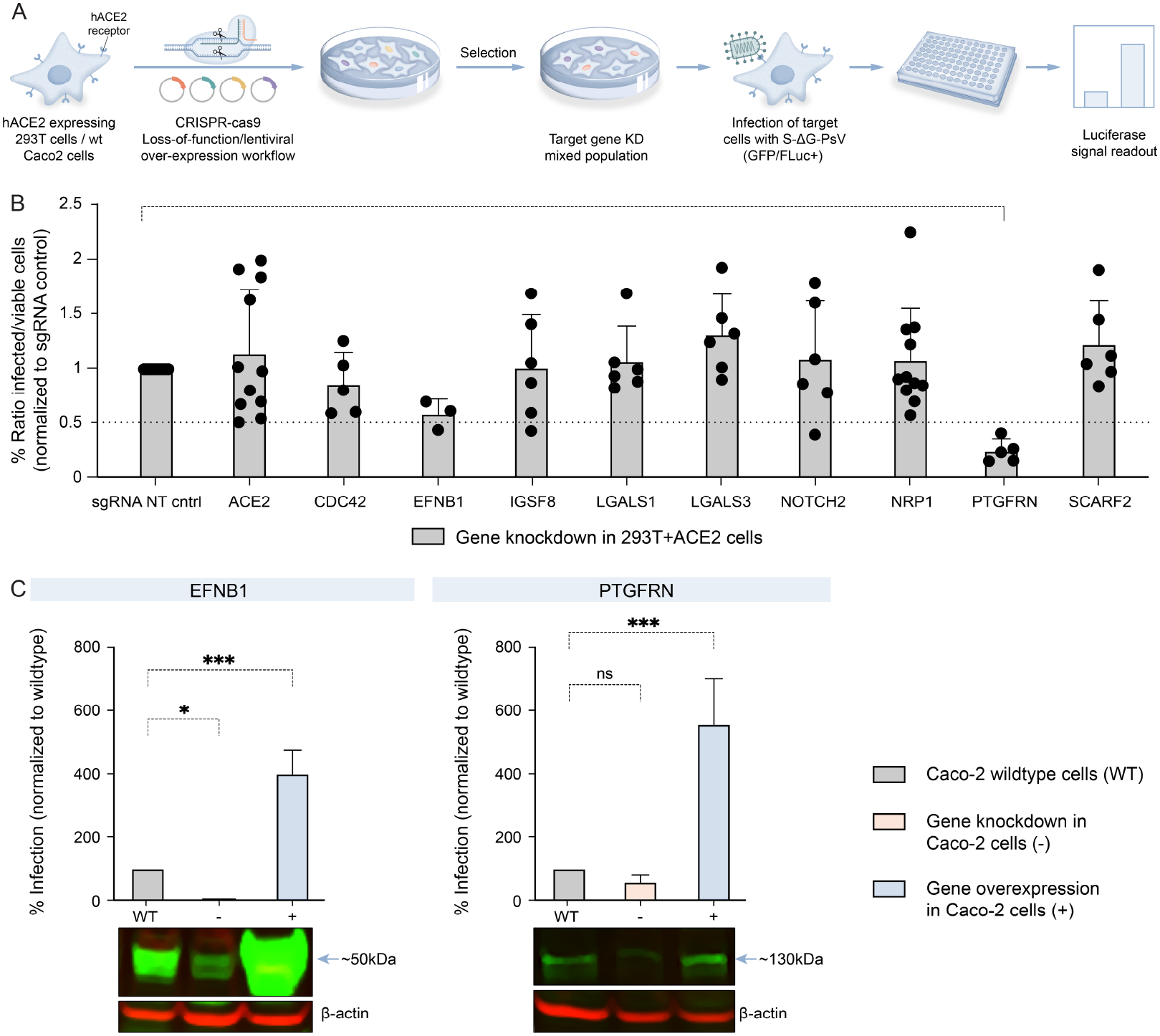
A. Screening and target validation workflow in pseudotype model. B. Effect of SARS CoV-2 Spike-typed pseudovirus infection on 293T+ACE2 knockdown cells for 10 HC host targets read by luciferase assay (experimental n = ≥ 3; one-way ANOVA with Dunnett’s test; mean ± s.d.) C. Effect of SARS CoV-2 Spike typed pseudovirus infection on loss and gain of function of EFNB1 and PTGFRN in Caco-2 cells analyzed by luciferase assay (experimental n= 3; one-way ANOVA with Dunnett’s test; mean ± s.d.) after confirming the gene knockdown and overexpression by western blot. All measurements are normalized to the controls for each assay.

To further confirm the role of PTGFRN and EFNB1 in SARS-CoV-2 entry, we either depleted these proteins or stably overexpressed them in Caco-2 cells and examined the ability of these cells to support SARS-CoV-2 pseudoparticle entry. Since we did not select for single-cell clones following CRISPR/Cas9-mediated genetic perturbation, we could only obtain partial depletion of proteins. The immunoblot analysis showed a depletion of 73.5% and 68.4% for PTGFRN and EFNB1, respectively (**Figure 3c & S3b**). Similarly, overexpression of PTGFRN and EFNB1 increased the cellular abundance of these proteins by 18.3% and 940.4%, respectively (**Figure 3c & S3b**). While PTGFRN depletion did not cause a significant change in pseudoparticle entry (**Table S3**), its overexpression increased the entry by 5.5 fold (p-value = 0.0005) (**Figure 3c**). On the other hand, EFNB1 overexpression significantly increased pseudoparticle entry (3-fold, p-value = 0.0001) in Caco-2 cells (Figure 3c), whereas its depletion caused a significant decrease in entry (16-fold, p-value = 0.0357) (**Figure 3c**, **Table S3**).

### PTGFRN and EFNB1 facilitate infection of authentic SARS-CoV-2

Next, we tested the role of PTGFRN and EFNB1 in SARS-CoV-2 entry by using authentic virus particles. Unfortunately, our Caco-2 cells showed extremely low susceptibility to SARS-CoV-2, precluding their use for defining minor changes in infection efficiency. As an alternative, we used HK2 cells, (a proximal tubular cell line derived from human kidney (Ryan et al., 1994), which, our previous work showed, express detectable levels of endogenous ACE2 both on the cell surface and intracellularly and support efficient SARS-CoV-2 infection (Chen *et al*., 2021). We knocked down PTGFRN and EFNB1 in HK2 cells, infected the cells with an mNeonGreen reporter SARS-CoV-2 virus (SARS-CoV-2-mNG) at a multiplicity of infection (MOI) of 0.5 or 3.0, and quantified the fluorescent signal of infected cells by flow cytometry at 12-hour and 24-hour post infection (h.p.i.) (**Figure 4a**).

**Figure 4.**
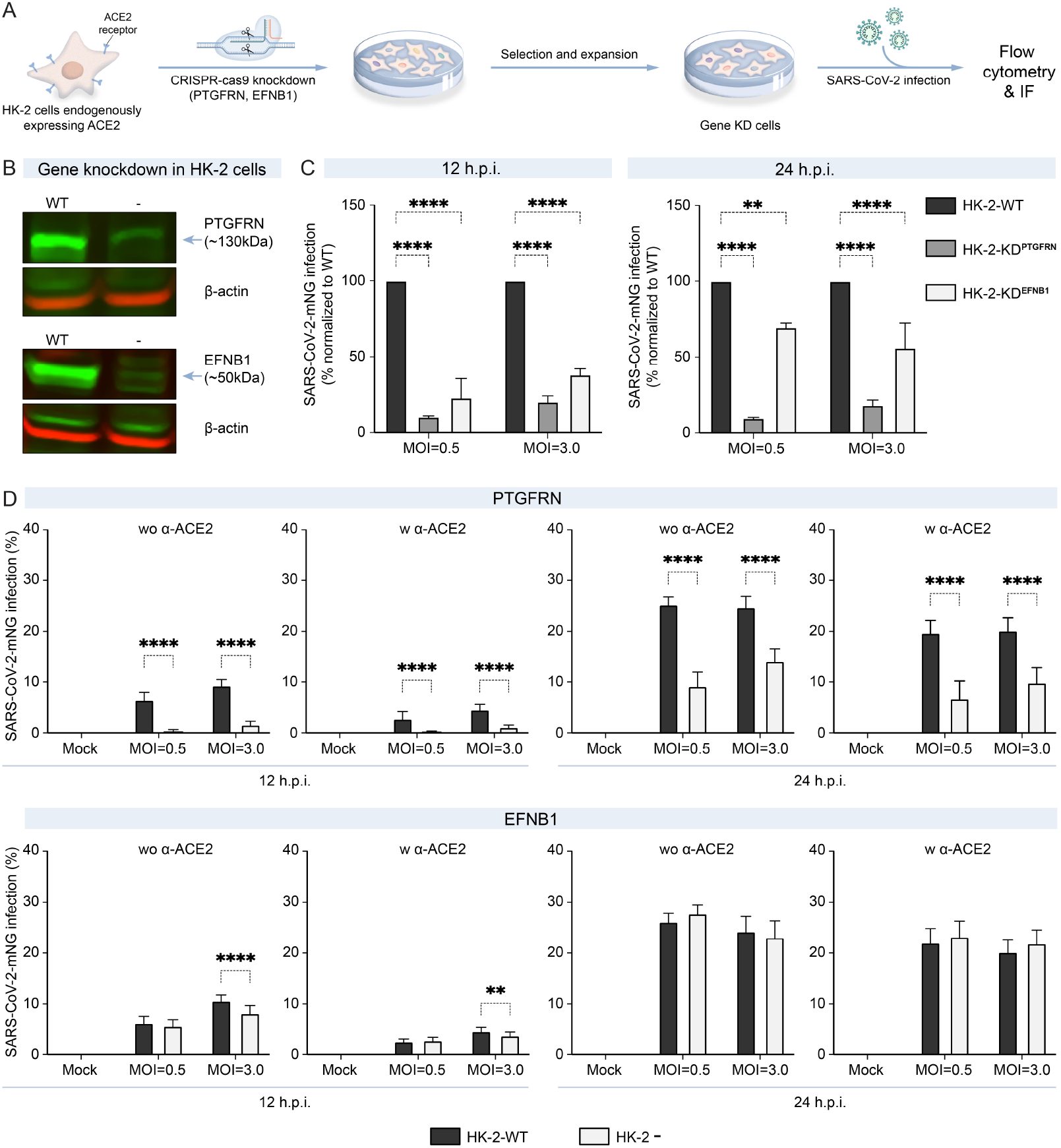
A. Target validation workflow in authentic live virus infection model. B. PTGFRN and EFNB1 expression knockdown in HK-2 cells C. Percentage of infected cell population after normalization upon knockdown of PTGFRN and EFNB1 in HK-2 cells at different multiplicity of infection (0.5 and 3.0) and time points (12 h.p.i. and 24 h.p.i.) of infection via flow cytometry. D. Bar plots showing percentage of SARS CoV-2 infection in wildtype (WT), PTGFRN and EFNB1 knockdown HK-2 cells at different multiplicity of infection (0.5 and 3.0) and time points of infection (12 h.p.i. and 24 h.p.i.) in presence and absence of ACE2-blockade analyzed from immunofluorescence (IF) imaging (technical replicate n = 10; two-way ANOVA with Šídák’s test; mean ± s.d.). All measurements are normalized to the controls for each assay.

At 12 h.p.i., PTGFRN and EFNB1 knockdown significantly reduced (~80-90%; *p*-value = <0.0001) viral infection (**Table S4**) relative to control cells (**Figure 4c** & **S4b**). At 24 h.p.i., a near complete (80-90%; *p*-value = <0.0001) decrease in infection was again seen for PTGFRN KD, whereas the infection blockade was partially alleviated (30-40% reduction in infection, *p*-value =<0.0001) for EFNB1 KD (**Figure 4c** & **S4b**).

To elucidate whether PTGFRN and EFNB1 work in concert with ACE2, we treated PTGFRN KD and EFNB1 KD HK-2 cells with an excess of anti-ACE2 antibody one hour prior to SARS-CoV-2 exposure (**see Methods section**), and measured potential change in infection relative to untreated cells. The IC_50_ concentration of anti-ACE2 antibody was determined empirically (**Figure S4c**). As expected, antibody-mediated blockade of ACE2 significantly reduced virus infection by ~50% when compared to untreated cells (**Table S5**). Notably, PTGFRN knockdown led to an additive reduction in infection in anti-ACE2 treated cells (**Figure 4d**), consistent with a possible role as an entry regulator in both ACE2 expressing and deficient cells. On the other hand, despite reproducing a similar trend of reducing infection preferentially at 12 h.p.i., the EFNB1 knockdown did not cause any further additive change in anti-ACE2 treated cells, consistent with its weaker effect on blocking infection at a later stage (**Table S5**).

## Discussion

Spike protein is a pivotal target for the development of therapeutic antibodies, entry inhibitors and vaccines as effective countermeasures for COVID-19. Since this viral protein is responsible for the initial SARS-CoV-2-host interactions and serves as the opening line of attack, we designed our study focusing on mapping its proximal membrane protein interactors on the host cell surface, aiming to contribute new co-receptor targets towards more lasting clinical measures.

Our viramapping datasets establish PTGFRN as a conserved target for the SARS virus family, suggesting it as a possible broad range therapeutic target for abrogating coronavirus infections. PTGFRN is known to physically interact with the glycoprotein binding partner of tetraspanins CD9 (Tspan29) and CD81 (Tspan28) that engage the actin cytoskeleton to control cell motility and polarity. Notably, CD81-PTGFRN interaction has previously been reported to both modulate HCV entry and influenza virus envelope formation (Feneant et al., 2014).Tspan29 is also known to promote MERS-CoV entry through condensation of its Spike protein and TEM-associated proteases on the cell surface (Earnest et al., 2017). Tspan expression has been implicated by Notch signaling (Ruiz-Garcia et al., 2016), which may explain the appearance of Notch-2 in our viramapping datasets. Evidence of interactions of PTGFRN and ITGB1 with galectins (LGALS1 and LGALS3) on the surface of retinal pigment epithelial cells (Obermann et al., 2017) may also justify the enrichment of all these proteins in our datasets. On the other hand, EFNB1, which is moderately expressed in vascular and kidney endothelial cells (Uhlen *et al*., 2015) and involved in IL-6 signaling during T cell-mediated viral immunity (Luo et al., 2011), enhanced virus entry, further implicating it as a second plausible therapeutic target.

Furthermore, our study demonstrates both the breadth of the Spike-host cell surface interactome and identifies additional, potentially transient interactors belonging to well-known membrane-tethered factors previously implicated in viral entry and host immune responses in other infection contexts. One notable highlight is the one-shot capture of the interactome comprising previously reported interactors in addition to novel ones in a single experiment, despite the fleeting and dynamic nature of some of these interactions. For example, the detection of integrin (ITGB1) and galectins (LGALS1 and LGALS3), recently reported to interact with SARS-CoV-2 Spike, in the Spike-ACE2 microenvironment, and the novel, validated SARS-CoV-2 host entry factors PTGFRN and EFNB1, identified uniquely in this study. The identification of these interacting host factors may explain the susceptibility of diverse human cells and tissues exhibiting low to undetectable ACE2 levels to SARS-CoV-2 infection (Chen *et al*., 2021; Li *et al*., 2020). The number of host factors in spatial proximity with the viral Spike protein also proves its promiscuous binding nature and motivates future studies aimed at uncovering their respective binding pockets on the trimeric Spike protein complex, as was recently documented for the NRP1, ganglioside and integrin-binding motifs on Spike (Lempp *et al*., 2021; Zhang *et al*., 2022).

In short, our study successfully deployed ViraMap technology using cell surface recognition targeting modalities to tease out both known and unknown virus-host protein interactions on the host outer cell membrane. We have established that this platform enables reliable identification of high confidence interactions at a high resolution with an unbiased affinity towards the closest neighbors of the viral Spike. Thus, we envision ViraMap, a photocatalyst-based proximity labeling platform, will emerge as a powerful next-generation interactome mapping approach that can be applied to unravel other clinically important virus-host interactomes. Another interesting potential extension of this application may involve attaching a photocatalyst onto other key SARS-CoV-2 viral effector proteins, such as the protease or replicase components that commandeer the intracellular host machinery, to identify their respective interactions and functional targets within the cytosolic environment.

## Supporting information

ViraMap Supplementary Information

## Acknowledgments

We thank Stefan Pölhman lab (University Göttingen, Göttingen, Germany) for pCG1-SARS-CoV-2-Spike and pVSV-ΔG-GFP-Fluc expression plasmids and Pei-Yong Shi lab (UTMB, Texas, USA) for the SARS-CoV-2-mNG reporter strain. We thank Avik Basu for advice interpreting the proteomic/mass spectrometry data, and Michal Depa for support with the image analysis. We also thank Da Yuan Chen for experimental support in BSL3 facilities.

We thank the Evans Center for Interdisciplinary Biomedical Research at Boston University School of Medicine for their support of the Affinity Research Collaborative on ‘Respiratory Viruses: A Focus on COVID-19’. We also thank Merck & Co. for supporting S.D. as principal research fellow during this project.

## Author Contributions

O.O.F., R.C.O. and A.E. co-conceived the study; O.O.F., R.C.O., M. S., and A.E. supervised the project; S.D., T.R.-R., O.O.F., R.C.O and M.S. designed methodology; S.D., A.H.T., T.R.-R., K.A.R., N.K., T.J.B., J.M.B., C.H.W., O.O.F and R.C.O executed experiments; S.D., T.R.-R., T.J.B., C.H.W., O.O.F., R.C.O. and M.S. analyzed data; D.J.H., K.A.V. and E.C.H. provided insight and direction for project design; S.D. prepared the original manuscript draft, T.R.-R., O.O.F., R.C.O., M.S. and A.E. reviewed and edited the manuscript.

## Materials & Correspondence

Correspondence and requests for materials should be addressed to lead contact, Andrew Emili

## Data Availability

The mass spectrometry proteomics data have been deposited to the MassIVE (ftp://MSV000089520@massive.ucsd.edu). All other data generated or analyzed during this study are included in this published article and its Supporting Information.

## RESOURCE AVAILABILITY

### Lead Contact

Requests for further information ad requests for reagents and resources should be directed to and will be fulfilled by Lead Contact, Andrew Emili

### Materials Availability

All unique reagents generated in this study are available from the Lead Contact without restriction for non-commercial applications, or with a completed Materials Transfer Agreement if there is potential for commercial applications; requestors are required to handle all shipping and customs costs.

### Data and Code Availability

MS data is available via MassIVE (ftp://MSV000089520@massive.ucsd.edu)

## EXPERIMENTAL MODEL AND SUBJECT DETAILS

### HEK-293T Human Embryonic Kidney Cells

HEK-293T cells were cultured under standard conditions in Dulbecco’s Modified Eagle Medium (DMEM) supplemented with 10% Fetal Bovine Serum and 1% Penicillin-Streptomycin at 37 °C at 5% CO_2_. Cells were passaged every 2 to 3 days (80-90% confluency) for 25-30 passages.

### Caco-2 Human Colon Carcinoma Cells

Caco-2 cells were cultured under standard conditions in Dulbecco’s Modified Eagle Medium (DMEM) (Gibco) supplemented with 10% Fetal Bovine Serum (Gibco) and 1% Penicillin-Streptomycin (Gibco) at 37 °C at 5% CO_2_. Cells were passaged every 2 to 3 days (80-90% confluency) for 25-30 passages.

### HK-2 Human Kidney Cells

HK-2 cells were cultured under standard conditions in Dulbecco’s Modified Eagle Medium (DMEM) (Gibco) supplemented with 10% FBS and 1% Penicillin-Streptomycin at 37 °C at 5% CO2. Cells were passaged every 2 to 3 days (80-90% confluency) for 25-30 passages.

## METHODS

### Preparation of IrPC labeled Spike protein, anti-His and anti-ACE2 antibody

200μL 1mg/mL SARS-CoV-2 Spike protein trimer (Acrobiosystems: SPN-C52H9), SARS-CoV-1 Spike trimer (Acrobiosystems: SPN-S52H5), SARS-CoV-2 Spike D614G (Acrobiosystems: SPN-C52H3), anti-His (Millipore: 05-949, clone HIS.H8) or Goat anti-Mouse IgG (Millipore: AP124) dissolved/diluted in PBS was transferred to an Eppendorf tube followed by addition of 25μL of 1M NaHCO3. 1.5 μL of 100mM C3-azide NHS ester (C3 linker, Enamine: EN300-265680) was added to the reaction vial and incubated for 2 hours at room temperature. The sample was then desalted using a Zeba spin column (ThermoFisher: 89891). The de-salted azide labeled protein was then labeled with IrPC alkyne (150μM final concentration) using the click-chemistry kit (ThermoFisher: C10276). The sample was incubated for 30 minutes followed by desalting on a Zeba spin column (MWCO 40kDa). Protein concentration was then determined by the BCA assay and diluted to a final concentration of 1mg/mL. The conjugation efficiency was determined using a standard conjugation evaluation protocol. Specifically, protein concentration is determined using the BCA assay with known amounts of BSA as a standard. The photocatalyst concentration was determined by measuring absorbance at 350nm and comparing to known amounts of free photocatalyst. The concentration of photocatalyst is divided by the concentration of protein to get the ratio.

### Flow cytometry analysis for Spike targeted viramapping on live cells

HEK293T+ACE2 cells (BPS Biosciences: 79951) were grown to confluency and detached by addition of Accutase (1min at 37°C). Cells were gently detached by pipetting and transferred to an Eppendorf tube. Cells were pelleted and resuspended at 2.5 million cells/mL in 0.5 mL cold PBS and incubated with 5μg IrPC labeled Spike protein or IgG and incubated for 60 min at 4C. For experiments involving ACE2 competition, 25 ug of ACE-Fc (Acrobiosystems: AC2-H5255) was added with the IrPC labeled Spike protein. After incubation, the cells were washed once with 1ml cold PBS and resuspended in 1mL cold PBS containing 250μM biotin-diazirine (prepared as described previously (Geri *et al*., 2020)) and incubated for 3min under blue light 100% intensity. The cells were washed 1× with 1ml cold PBS and resuspended in 500μl PBS containing PE Streptavidin (BD 554061: 1:200 dilution) and Zombie Violet live dead stain (BioLegend 423113: 1:500 dilution). The cells were washed 0.5mL 1× cold PBS and analyzed by flow cytometry for biotinylation.

### Confocal microscopy imaging of Spike targeted viramapping on live cells

For photocatalyst labelling method, 500,000 cells/ml HEK293+ACE2 cells were plated into a ibidi microscopy plates and incubated for 24 hours at 37°C in complete media after pre-coating with poly-L-lysine for 30min at 37°C. After 24-hour incubation, the cells were washed once with 1mL cold complete DMEM media and incubated in cold DMEM containing 10μg/ml Spike-IrPC or IgG-IrPC for 30 minutes at 4C. The media was then removed, and the plates were washed twice with 1mL complete DMEM media. 0.5mL of cold PBS containing 250μM biotin-diazirine (prepared as described previously (Geri *et al*., 2020)) was then added to the cell plates. The cells were then irradiated for 3min in the photoreactor for at 100% intensity. The cells were then washed twice with PBS and fixed with 6% paraformaldehyde (PFA, Electron Microscopy Sciences: 15710) and 0.2% glutaraldehyde (Sigma-Aldrich: G5882-10X10ML) that were prepared in 1× DPBS and added gently at equal ratios per dish (final concentration of 3% PFA and 0.1% glutaraldehyde in a total volume of 400 μL) and incubated for 10 min at 4°C. The dishes were washed 3× in Stain Buffer (BD Biosciences: 554656) and incubated overnight in 1 mL of Stain Buffer at 4°C. The following day, samples were stained with Alexa Fluor 488 Streptavidin (BioLegend: 405235) at a 1:200 dilution in 400 μL of Stain Buffer and incubated overnight at 4°C. The samples were washed 1× with 1 mL of Stain Buffer and Hoechst DNA dye (Cayman Chemical Company: 600332) was added at a 1:10,000 dilution in 400 μL of Stain Buffer per dish and incubated while protected from light for 10 min at room temperature. The dishes were washed 2× in Stain Buffer and fixed with 400 μL of a 3% PFA and 0.1% Glutaraldehyde solution in 1× DPBS for 5 min at room temperature, washed 2× in 1 mL of Stain Buffer, and imaged using a Zeiss LSM800 inverted, confocal microscope using a 63X oil immersion objective.

### Photolabeling on live cells for quantitative LC-MS/MS analysis

HEK293+ACE2 cells (BPS Biosciences: 79951) were grown in 15cm culture dishes to confluency. Cells were detached using 3mL cold Accutase added to each plate and incubated for 1min at 37°C. 5mL of media was added to each plate and cells were detached by pipetting multiple times. Cells were collected into a 50mL conical vial and centrifuged to pellet the cells (5min, 800×g, 5min). The media was removed and the cells were resuspended in cold PBS in each falcon tube and combined into a single falcon tube and pelleted by centrifugation (5min, 800×g at 4°C). After centrifugation, the supernatant was removed and cells were resuspended in cold PBS at 20 million cells/mL and then distributed into 1.5 mL microcentrifuge tubes (Eppendorf Protein LoBind tubes (Z666505-100EA) at 1 mL each. The tubes were pelleted by centrifugation and resuspended in 1mL cold PBS. For negative control samples, 20μg of goat-anti mouse-IrPC (Millipore: AP124) was added. 20μg of each of three Spike-IrPC variants, or anti-ACE2 antibody (R & D systems: MAB9332) were added. Both control and test samples were incubated on a nutator at 4°C for 60 min followed by 2× 1mL cold PBS washes (cells centrifuged at 800×g for 4 min at 4°C to pellet cells). For anti-ACE2 labeling, 20μg of goat-anti mouse-IrPC was added to each sample and incubated at 4°C for 30 min followed by 2× 1mL cold PBS washes (cells centrifuged at 800×g for 4 minutes at 4°C to pellet cells). For labeling with anti-His tag IrPC, the anti-His tag IrPC conjugate was added to the Spike-labeled cells and incubated at 4°C for 30 min followed by 2× 1mL cold PBS washes (cells centrifuged at 800×g for 4 min at 4°C to pellet cells). After coating the cells with IrPC-labeled Spike or antibody, the cells were resuspended in 1mL of cold PBS containing 250μM biotin diazirine probe (prepared as described previously (Geri *et al*., 2020)). The samples were placed in the BPR200 for 3min and irradiated with blue light (450 nm) at 100% intensity. Cells were then washed twice with cold PBS by spinning at 800xg for 4 minutes. 1mL of membrane permeabilization buffer (MEM-PER plus membrane fractionation kit Thermo: 89842 with protease inhibitors) was added to the cells and incubated for 20min on ice. After incubation, the cells were then spun at 16,000×g for 15min at 4°C. The supernatant was removed and discarded. For the remaining membrane fraction pellet, 300μL of lysis buffer (RIPA with 1% SDS and 1× protease inhibitor cocktail) was added to Eppendorf tube and the samples were sonicated to break-up the membrane pellet (1×5s and power 6). The samples were boiled for 5min at 95°C. The samples were then diluted to 1.3mL with RIPA and sonicated till lysate viscosity was consistent with aqueous buffer (2×5s power 7). The protein concentration was measured by BCA and the samples were stored at −80°C till streptavidin bead enrichment. For bead enrichment, 250uL of streptavidin magnetic beads (Thermo Fisher: 88817) were washed with 2×1mL RIPA. 2mg of each sample was then added to the beads and incubated for 3 hours at RT on a rotisserie. After incubation, the beads were washed thrice each using 1mL 1%SDS in PBS (5min incubation in between), 1mL 1M NaCl in PBS (5min incubation in between), 1mL 10% EtOH in PBS (5min incubation in between) and 1mL RIPA. After the RIPA wash the beads were treated with 30μl of 4× loading buffer (Bio-Rad: 161-0747) containing 20mM DTT and 25mM biotin (no BME) was added to the beads in each tube and boiled at 95°C for 10min. The beads were pelleted using a magnetic separator and the supernatant was removed. The supernatants were then stored at −80°C till MS/MS proteomic analysis (IQ proteomics).

### LC-MS/MS workflow (TMT workflow and instrument parameters) and quantitative peptide search

Quantitative proteomic sample preparation and analysis was performed at IQ Proteomics, Cambridge, MA. Proteins were reduced with 20 mM dithiothreitol (DTT) for 1 hour at room temperature. Cysteine residues were alkylated with iodoacetamide (60 mM) for 1hour in the dark and quenched with DTT (40 mM). Proteins were extracted by methanol-chloroform precipitation and digested with 1 μg of trypsin (Promega) in 100 mM EPPS (pH 8.0) for 4 hours at 37 °C. Each of the tryptic peptide samples were labeled with 400 μg of Tandem Mass Tag (TMT; Pierce) isobaric reagents for 2 hours at room temperature. All samples were quenched with hydroxylamine (0.5%), acidified with TFA (2%), pooled, and dried by speedvac evaporation. Pooled TMT labeled peptides were fractionated using the high pH reverse-phase peptide fractionation kit (Pierce) into 3 fractions (20%, 25%, and 50% acetonitrile in 0.1% triethylamine) and desalted with Empore-C18 (3M) in-house packed StageTips prior to analysis by mass spectrometry. All mass spectra were acquired on an Orbitrap Fusion Lumos coupled to an EASY nanoLC-1000 (or nanoLC-1200) (Thermo Fisher) liquid chromatography system. Approximately 2 μg of peptides were loaded on a 75 μm capillary column packed in-house with Sepax GP-C18 resin (1.8 μm, 150 Å, Sepax) to a final length of 35 cm. Peptides were separated using a 110-minute linear gradient from 8% to 28% acetonitrile in 0.1% formic acid. The mass spectrometer was operated in a data dependent mode. The scan sequence began with FTMS1 spectra (resolution = 120,000; mass range of 350–1400 m/z; max injection time of 50 ms; AGC target of 1e6; dynamic exclusion for 60 seconds with a +/− 10 ppm window). The ten most intense precursor ions were selected for MS2 analysis via collisional-induced dissociation (CID) in the ion trap (normalized collision energy (NCE) = 35; max injection time = 100ms; isolation window of 0.7 Da; AGC target of 1.5e4). Following MS2 acquisition, a synchronous-precursor selection (SPS) MS3 method was enabled to select eight MS2 product ions for high energy collisional-induced dissociation (HCD) with analysis in the Orbitrap (NCE = 55; resolution = 50,000; max injection time = 86 ms; AGC target of 1.4e5; isolation window at 1.2 Da for +2 m/z, 1.0 Da for +3 m/z or 0.8 Da for +4 to +6 m/z). All mass spectra were converted to mzXML using a modified version of ReAdW.exe. MS/MS spectra were searched against a concatenated 2018 human Uniprot protein database containing common contaminants (forward + reverse sequences) using the SEQUEST algorithm (Eng et al., 1994). Database search criteria were as follows: fully tryptic with two missed cleavages; a precursor mass tolerance of 50 ppm and a fragment ion tolerance of 1 Da; oxidation of methionine (15.9949 Da) was set as differential modifications. Static modifications were carboxyamidomethylation of cysteines (57.0214) and TMT on lysines and N-termini of peptides (229.1629). Peptide-spectrum matches were filtered using linear discriminant analysis (Huttlin et al., 2010) and adjusted to a 1% peptide false discovery rate (FDR) (Elias and Gygi, 2007).

### Data analysis for MS raw data

All bioinformatic analysis of LC-MS/MS data including peptide level abundance, FDR corrected p-values and fold-change generation was performed in the R statistical computing environment as previously described (Geri *et al*., 2020). Peptide level abundance data of each protein was normalized to the summed total abundance for each sample. These totals were then averaged, and each normalized protein abundance value is multiplied by this average to rescale abundance data. Peptide level data is then merged to protein level data by taking the median of all peptides corresponding to a protein. Proteins were then filtered to remove any known contaminants identified from the database search. The enriched proteins were visualized by volcano plots generated in the manuscript by using GraphPad Prism 9.0. Proteins were colored and labeled based on whether they fell above or below the log_2_-fold cutoff threshold (≥1.5) and were statistically significant (-log_10_.Value ≥ 2.0). To identify a reliable and comprehensive list of biologically relevant protein interactions based on their abundance, reproducibility and specificity within and across experiments, the reporter ion intensities for all identified proteins from the MS-raw data were then subjected to high confidence interaction scoring pipeline using both Mass spectrometry interaction STatistics (MiST) (Jager *et al*., 2011; Verschueren *et al*., 2015) and Significance Analysis of INTeractome express (SAINT express) (v.3.6.3) (Teo *et al*., 2014) as previously described. All protein interactions that had a MiST score ≥ 0.8 and a SAINTexpress Bayesian false-discovery rate (BFDR) ≤ 0.3 for triplicate experiments and ≤ 0.01 for single experiments were chosen for network analysis using Cytoscape 3.9.1 (Morris et al., 2014). An organic layout was applied to the network with nodes representing the proteins, and edges representing the average of MiST score and BFDR for each interaction in the network. The nodes were grouped based on the interaction overlap with different types of viramap nodes. Cytoscape plugin ClueGO was used for the Gene Ontology (GO) enrichment analysis of the output list of 18 HC shared host factors (Bindea et al., 2009). GO database of GO_BiologicalProcess-EBI-UniProt-GOA-ACAP-ARAP_13.05.2021_00h00 was selected. GO terms were clustered based on shared gene members using kappa statistics to link the terms in the network. Similar GO terms were represented as nodes of the same color and the edges were the quantitative representation of shared gene membership (kappa score). Enrichment significance of the terms was represented by the size of the nodes. The p value representing the chance of observing n number of genes in a gene list annotated to a specific term was set at <0.05 keeping the default ClueGO features.

### Cell culture, plasmid cloning and generating stable cell lines

Human embryonic kidney HEK293T cells (ATCC: CRL-3216), human epithelial colorectal adenocarcinoma Caco-2 cells (ATCC: HTB-37), and human epithelial kidney papilloma HK-2 cells (ATCC: CRL-2190) were maintained at 37°C and 5% CO_2_ in DMEM containing 10% FBS. The human NRP1 (HG10011-G), EFNB1 (HG10894-M), PTGFRN (HG20704-U) were purchased from SinoBiological and cloned into the lentiviral pLOC plasmid to obtain pLOC-EFNB1 and pLOC-PTGFRN, respectively. The pLOC_hACE2_PuroR plasmid containing the human ACE2 has previously been described (Chen *et al*., 2021). The cas9-carrying lentiviral vector spCas9-BlastR (plasmid no. 71489) was obtained from Addgene. pCG1-SARS-CoV-2_S plasmid bearing the codon-optimized form of the SARS-CoV-2 spike gene, and pVSV-ΔG-GFP-Fluc plasmid, carrying the replication-competent vesicular stomatitis virus (VSV) genome with the viral glycoprotein (G) gene being replaced with GFP and firefly luciferase, were obtained from Stefan Pöhlmann (Hoffmann *et al.*, 2020). Stable cells expressing Cas9, ACE2, EFNB1 and PTGFRN were generated by transducing them with lentivirus particles followed by selection with appropriate selection markers.

### Generating CRISPR-Cas9 mediated gene knockdown cells

The predesigned Dharmacon™ Edit-R™ lentiviral sgRNA plasmids (set of 3) for the selected list of 10 genes were purchased from Horizon Discovery (https://horizondiscovery.com). Lentivirus particles were packaged in Lenti-X cells (ThermoFisher: NC9834960). Reverse transduction was carried out to generate the knock-down cells. Briefly, for each well in a 6-well plate, 150,000 cells mixed with appropriate sgRNA lentiviral stock and polybrene were plated and incubated at 37°C and 5% CO_2_. After 48 hours, the culture medium was replaced with fresh medium containing 5 μg/ml Puromycin. Cells were selected for 48 hours and used for cell viability or infection assays.

### Pseudovirus preparation and entry assay

To generate SARS-CoV-2 pseudoparticles, HEK293T cells were seeded into 6-well plates at a density of 600,000 cells per well. The following day, the cells were transfected with pCG1-SARS-CoV-2_S plasmid using X-tremeGENE™ 9 DNA transfection reagent (Millipore-Sigma: 06365779001) (ratio 3:1 μl/μg DNA). Infection with GFP/Firefly luciferase-expressing ΔG-Vesicular stomatitis virus (VSV) particles was carried out 20-hour post-transfection at MOI 3.0. Cells were gently washed three times 2-hour post-infection and incubated in DMEM containing 1:1000 dilution of anti-VSV-G antibody (Kerafast: 8G5F11) to eliminate residual ΔG-VSV particles at 37°C. Culture media were collected after 24 hours and centrifuged at 1000 × g for 5 min to discard the cell debris. Supernatants with pseudoviral particles were filtered through 0.45μm using syringe into fresh conical tubes. Stocks were further concentrated by 20% sucrose gradient ultracentrifugation.

For entry assays, each well of a 96-well plate were seeded with 20,000 cells one day prior to infection. For flow-cytometry assay, each well of a 24-well plate was seeded with 150,000 cells one day prior to infection. The following day, cells were checked for 60-70% confluency and infected with 1:3 dilution of the pseudoviral stock in Opti-MEM (ThermoFisher: 31985062). Cells were incubated in low serum condition for 24 hours at 37°C before both luciferase and flow cytometry assays.

### mNeonGreen-SARS-CoV-2 reporter virus stock preparation, anti-ACE2 antibody blocking and infection assay in BSL3

The recombinant SARS-CoV-2 expressing mNeonGreen (SARS-CoV-2-mNG) was obtained from Dr. Pei-Yong Shi at the University of Texas Medical Branch (UTMB). To generate working stock of the virus, Vero E6 cells were infected at an MOI of 0.01 for three days and the harvested cell culture medium was passed through a 0.45 μ filter. The virus stock was stored at −80°C in 1 ml aliquots. For the anti-ACE2 blocking assay, cells were treated for 2 hours prior to infection with the IC50 concentration of anti-ACE2 antibody (R&D Systems: AF933) determined for the cell lines used. For the infection assay, anti-ACE2 antibody solution was removed, and cells were infected with SARS-CoV-2-mNG in Opti-MEM for 1 hour at 37°C at an MOI of 0.5 or 3.0. After incubation, DMEM supplemented with 2% FBS was added to the cells and incubated at 37°C. Cells were harvested at 12 or 24 hours where indicated.

### Cell viability assay

Cell lysates were prepared as per the manufacturers protocol using the CellTiter-Glo® Luminescent Cell Viability Assay kit (Promega: G7570). Briefly, 48-hour post-perturbation (knockdown/overexpression), the culture medium was removed and replaced with equal volume of CellTitre-Glo® reagent, mixed well and incubated for 10 min at room temperature (RT) in dark. Luminiscence was measured in a Varioskan™ LUX multimode microplate reader (ThermoFisher). Percentage viability was calculated relative to parental control cells (100% viability).

### Luciferase assay for infected cells

Luciferase counts were measured for cells infected with Spike-pseudotyped VSV-GFP/Fluc particles using Luc-Pair™ Firefly Luciferase HS Assay Kit (GeneCopoeia: LF009). Briefly, 24-hour post-infection in gene-perturbed cells (knockdown/overexpression), the culture medium was removed and replaced with 1× lysis buffer (50μl per well of a 96-well plate). Cell suspensions were placed on a plate shaker at 4°C for 15 min at 400rpm to induce lysis. Cell lysates were centrifuged for 5 min at 1000× g to pellet cell debris. Supernatants were transferred to white opaque 96-well plate and mixed with 1× luciferase substrate reagent. Luciferase activity was recorded on Varioskan™ LUX multimode microplate reader (ThermoFisher). Percentage infection was calculated relative to parental control cells (100% infection).

### Flow cytometry for infected cells

Cells infected with SARS-CoV-2-mNG were harvested at the indicated timepoints, fixed in 10% formalin for 30 minutes, and washed once in 1× PBS. Data were acquired using a LSRII flow cytometer (BD) and analyzed with FlowJo version 10.1 software (Tree star). Infected cells were calculated as the percentage of NG positive cells per sample.

### IF imaging of infected cells and analysis

Virus-infected cells were fixed in 4% PFA for 30 min. The fixative was removed and the cell monolayer washed twice with 1× PBS. The cells were then permeabilized and incubated overnight at 4°C with anti-SARS-CoV-2 nucleocapsid antibody (Rockland: 200-401-A50; 1:2,000 dilution). The cells were then washed 5 times with 1× PBS and stained with Alexa Fluor 568-conjugated goat anti-rabbit secondary antibody (1:1,000 dilution) (Invitrogen: A11008) in the dark at room temperature for 1 hour and counterstained with 4′,6-diamidino-2-phenylindole (DAPI). Images were captured using an EVOS M5000 imaging system (ThermoFisher Scientific). For quantitative analysis of the fixed cell images, we used the MuviCyte live-cell imaging system (PerkinElmer, Waltham, MA). We acquired images of multiple microscopic fields per well using a 4× lens objective and counted the number of DAPI- and SARS-CoV-2 viral nucleoprotein (N)-positive cells. Composite confocal images were generated using a custom Python script. Images were processed and analyzed for cell counts by using CellProfiler 4.2 (McQuin et al., 2018). Percentage of DAPI-positive cells expressing the viral N protein was calculated for each image and plotted the mean ± standard deviation (SD) of multiple images (n=10) for each condition.

### Immunoblotting of gene-knockdown and overexpression cells

Proteins were extracted with 1× RIPA buffer containing 1× complete-mini protease inhibitor (Roche: 11836170001) and 1× phosphatase inhibitor cocktail (Roche: 04906837001). Samples were incubated on ice for 30 min and centrifuged at 12,000 × g for 20 min at 4°C. The supernatants were transferred to new ice-cold Eppendorf tubes, and protein concentration was measured by the bicinchoninic acid (BCA) assay using the Pierce BCA protein assay kit (ThermoFisher Scientific: 23225). Equal amounts of protein were loaded on 4 to 12% SDS-PAGE gel and transferred onto nitrocellulose membrane. Anti-PTGFRN antibody (R&D Systems: MAB100431; 1:1000), anti-EFNB1 antibody (Invitrogen: 2D3E9; 1:1000), anti-NRP1 antibody (Abcam: EPR3113; 1:1000), mouse b-actin antibody (1:3000) and rabbit b-actin antibody (1:3000) were used for primary staining of the membranes in appropriate pairing, followed by staining with secondary antibodies (Li-Cor: 926-32212 and 926-68072) (1:20,000). The bands were visualized by scanning the membranes with the Li-Cor CLx infrared scanner.

### Statistical analysis

Experimental data from cellular and infection assays were analyzed using GraphPad Prism 9.0 and subjected to student’s t-test, one-way or two-way ANOVA, where appropriate, with representation from at least three independent experiments. P < 0.05 was considered significant.

## Titles of excel tables

**Table S1.** Protein proximity scoring results for all virmaps and all host prey proteins, showing MiST score and SAInt BFDR across experimental samples.

**Table S2.** Enrichment of GO Biological processes of 18 high-confidence (HC) host factors in SARS-CoV-2 Spike-ACE2 microenvironment.

**Table S3.** Pseudovirus infection assay and cell viability assay results for gene-perturbation in HEK293T+ACE2 cells and Caco-2 cells.

**Table S4.** Flow cytometry results for mNeonGreen reporter SARS-CoV-2 virus (SARS-CoV-2-mNG) infection in PTGFRN and EFNB1 knockdown HK-2 cells at different multiplicity of infection (0.5 and 3.0) and time points (12 h.p.i. and 24 h.p.i.).

**Table S5.** Cell counts after SARS CoV-2 infection in wildtype (WT), PTGFRN and EFNB1 knockdown HK-2 cells at different multiplicity of infection (0.5 and 3.0) and time points of infection (12 h.p.i. and 24 h.p.i.) in presence and absence of ACE2-blockade analyzed from immunofluorescence (IF) imaging.

## References

Amraei, R., Xia, C., Olejnik, J., White, M.R., Napoleon, M.A., Lotfollahzadeh, S., Hauser, B.M., Schmidt, A.G., Chitalia, V., Muhlberger, E., et al. (2022). Extracellular vimentin is an attachment factor that facilitates SARS-CoV-2 entry into human endothelial cells. Proc Natl Acad Sci U S A 119. 10.1073/pnas.2113874119.

Beauvillain, C., Meloni, F., Sirard, J.C., Blanchard, S., Jarry, U., Scotet, M., Magistrelli, G., Delneste, Y., Barnaba, V., and Jeannin, P. (2010). The scavenger receptors SRA-1 and SREC-I cooperate with TLR2 in the recognition of the hepatitis C virus non-structural protein 3 by dendritic cells. J Hepatol 52, 644–651. 10.1016/j.jhep.2009.11.031.

Bindea, G., Mlecnik, B., Hackl, H., Charoentong, P., Tosolini, M., Kirilovsky, A., Fridman, W.H., Pages, F., Trajanoski, Z., and Galon, J. (2009). ClueGO: a Cytoscape plug-in to decipher functionally grouped gene ontology and pathway annotation networks. Bioinformatics 25, 1091–1093. 10.1093/bioinformatics/btp101.

Bojkova, D., Klann, K., Koch, B., Widera, M., Krause, D., Ciesek, S., Cinatl, J., and Munch, C. (2020). Proteomics of SARS-CoV-2-infected host cells reveals therapy targets. Nature 583, 469–472. 10.1038/s41586-020-2332-7.

Bowden, T.A., Aricescu, A.R., Gilbert, R.J., Grimes, J.M., Jones, E.Y., and Stuart, D.I. (2008). Structural basis of Nipah and Hendra virus attachment to their cell-surface receptor ephrin-B2. Nat Struct Mol Biol 15, 567–572. 10.1038/nsmb.1435.

Burbage, M., Keppler, S.J., Gasparrini, F., Martinez-Martin, N., Gaya, M., Feest, C., Domart, M.C., Brakebusch, C., Collinson, L., Bruckbauer, A., and Batista, F.D. (2015). Cdc42 is a key regulator of B cell differentiation and is required for antiviral humoral immunity. J Exp Med 212, 53–72. 10.1084/jem.20141143.

Chen, D.Y., Khan, N., Close, B.J., Goel, R.K., Blum, B., Tavares, A.H., Kenney, D., Conway, H.L., Ewoldt, J.K., Chitalia, V.C., et al. (2021). SARS-CoV-2 Disrupts Proximal Elements in the JAK-STAT Pathway. J Virol 95, e0086221. 10.1128/JVI.00862-21.

Darling, T.K., and Lamb, T.J. (2019). Emerging Roles for Eph Receptors and Ephrin Ligands in Immunity. Front Immunol 10, 1473. 10.3389/fimmu.2019.01473.

Dermody, T.S., Kirchner, E., Guglielmi, K.M., and Stehle, T. (2009). Immunoglobulin superfamily virus receptors and the evolution of adaptive immunity. PLoS Pathog 5, e1000481. 10.1371/journal.ppat.1000481.

Earnest, J.T., Hantak, M.P., Li, K., McCray, P.B., Jr., Perlman, S., and Gallagher, T. (2017). The tetraspanin CD9 facilitates MERS-coronavirus entry by scaffolding host cell receptors and proteases. PLoS Pathog 13, e1006546. 10.1371/journal.ppat.1006546.

El Amri, M., Fitzgerald, U., and Schlosser, G. (2018). MARCKS and MARCKS-like proteins in development and regeneration. J Biomed Sci 25, 43. 10.1186/s12929-018-0445-1.

Elias, J.E., and Gygi, S.P. (2007). Target-decoy search strategy for increased confidence in large-scale protein identifications by mass spectrometry. Nat Methods 4, 207–214. 10.1038/nmeth1019.

Eng, J.K., McCormack, A.L., and Yates, J.R. (1994). An approach to correlate tandem mass spectral data of peptides with amino acid sequences in a protein database. J Am Soc Mass Spectrom 5, 976–989. 10.1016/1044-0305(94)80016-2.

Feneant, L., Levy, S., and Cocquerel, L. (2014). CD81 and hepatitis C virus (HCV) infection. Viruses 6, 535–572. 10.3390/v6020535.

Geri, J.B., Oakley, J.V., Reyes-Robles, T., Wang, T., McCarver, S.J., White, C.H., Rodriguez-Rivera, F.P., Parker, D.L., Jr., Hett, E.C., Fadeyi, O.O., et al. (2020). Microenvironment mapping via Dexter energy transfer on immune cells. Science 367, 1091–1097. 10.1126/science.aay4106.

Gordon, D.E., Jang, G.M., Bouhaddou, M., Xu, J., Obernier, K., White, K.M., O’Meara, M.J., Rezelj, V.V., Guo, J.Z., Swaney, D.L., et al. (2020). A SARS-CoV-2 protein interaction map reveals targets for drug repurposing. Nature 583, 459–468. 10.1038/s41586-020-2286-9.

Hantak, M.P., Qing, E., Earnest, J.T., and Gallagher, T. (2019). Tetraspanins: Architects of Viral Entry and Exit Platforms. J Virol 93. 10.1128/JVI.01429-17.

Hoffmann, M., Kleine-Weber, H., Schroeder, S., Kruger, N., Herrler, T., Erichsen, S., Schiergens, T.S., Herrler, G., Wu, N.H., Nitsche, A., et al. (2020). SARS-CoV-2 Cell Entry Depends on ACE2 and TMPRSS2 and Is Blocked by a Clinically Proven Protease Inhibitor. Cell 181, 271–280 e278. 10.1016/j.cell.2020.02.052.

Huttlin, E.L., Jedrychowski, M.P., Elias, J.E., Goswami, T., Rad, R., Beausoleil, S.A., Villen, J., Haas, W., Sowa, M.E., and Gygi, S.P. (2010). A tissue-specific atlas of mouse protein phosphorylation and expression. Cell 143, 1174–1189. 10.1016/j.cell.2010.12.001.

Jager, S., Cimermancic, P., Gulbahce, N., Johnson, J.R., McGovern, K.E., Clarke, S.C., Shales, M., Mercenne, G., Pache, L., Li, K., et al. (2011). Global landscape of HIV-human protein complexes. Nature 481, 365–370. 10.1038/nature10719.

Lempp, F.A., Soriaga, L.B., Montiel-Ruiz, M., Benigni, F., Noack, J., Park, Y.J., Bianchi, S., Walls, A.C., Bowen, J.E., Zhou, J., et al. (2021). Lectins enhance SARS-CoV-2 infection and influence neutralizing antibodies. Nature 598, 342–347. 10.1038/s41586-021-03925-1.

Li, M.Y., Li, L., Zhang, Y., and Wang, X.S. (2020). Expression of the SARS-CoV-2 cell receptor gene ACE2 in a wide variety of human tissues. Infect Dis Poverty 9, 45. 10.1186/s40249-020-00662-x.

Lim, P.S., Sutton, C.R., and Rao, S. (2015). Protein kinase C in the immune system: from signalling to chromatin regulation. Immunology 146, 508–522. 10.1111/imm.12510.

Liu, S., Zhao, W., Li, X., Zhang, Gao, Y., Peng, Q., Du, C., and Jiang, N. (2021). AGTRAP Is a Prognostic Biomarker Correlated With Immune Infiltration in Hepatocellular Carcinoma. Front Oncol 11, 713017. 10.3389/fonc.2021.713017.

Luo, H., Charpentier, T., Wang, X., Qi, S., Han, B., Wu, T., Terra, R., Lamarre, A., and Wu, J. (2011). Efnb1 and Efnb2 proteins regulate thymocyte development, peripheral T cell differentiation, and antiviral immune responses and are essential for interleukin-6 (IL-6) signaling. J Biol Chem 286, 41135–41152. 10.1074/jbc.M111.302596.

Lyu, J., Imachi, H., Fukunaga, K., Yoshimoto, T., Zhang, H., and Murao, K. (2015). Roles of lipoprotein receptors in the entry of hepatitis C virus. World J Hepatol 7, 2535–2542. 10.4254/wjh.v7.i24.2535.

Maginnis, M.S. (2018). Virus-Receptor Interactions: The Key to Cellular Invasion. J Mol Biol 430, 2590–2611. 10.1016/j.jmb.2018.06.024.

Martin-Vilchez, S., Whitmore, L., Asmussen, H., Zareno, J., Horwitz, R., and Newell-Litwa, K. (2017). RhoGTPase Regulators Orchestrate Distinct Stages of Synaptic Development. PLoS One 12, e0170464. 10.1371/journal.pone.0170464.

Martinez, W.M., and Spear, P.G. (2001). Structural features of nectin-2 (HveB) required for herpes simplex virus entry. J Virol 75, 11185–11195. 10.1128/JVI.75.22.11185-11195.2001.

May, D.G., Martin-Sancho, L., Anschau, V., Liu, S., Chrisopulos, R.J., Scott, K.L., Halfmann, C.T., Diaz Pena, R., Pratt, D., Campos, A.R., and Roux, K.J. (2022). A BioID-Derived Proximity Interactome for SARS-CoV-2 Proteins. Viruses 14. 10.3390/v14030611.

McQuin, C., Goodman, A., Chernyshev, V., Kamentsky, L., Cimini, B.A., Karhohs, K.W., Doan, M., Ding, L., Rafelski, S.M., Thirstrup, D., et al. (2018). CellProfiler 3.0: Next-generation image processing for biology. PLoS Biol 16, e2005970. 10.1371/journal.pbio.2005970.

Mellacheruvu, D., Wright, Z., Couzens, A.L., Lambert, J.P., St-Denis, N.A., Li, T., Miteva, Y.V., Hauri, S., Sardiu, M.E., Low, T.Y., et al. (2013). The CRAPome: a contaminant repository for affinity purification-mass spectrometry data. Nat Methods 10, 730–736. 10.1038/nmeth.2557.

Morris, J.H., Knudsen, G.M., Verschueren, E., Johnson, J.R., Cimermancic, P., Greninger, A.L., and Pico, A.R. (2014). Affinity purification-mass spectrometry and network analysis to understand protein-protein interactions. Nat Protoc 9, 2539–2554. 10.1038/nprot.2014.164.

Oakley, J.V., Buksh, B.F., Fernandez, D.F., Oblinsky, D.G., Seath, C.P., Geri, J.B., Scholes, G.D., and MacMillan, D.W.C. (2022). Radius measurement via super-resolution microscopy enables the development of a variable radii proximity labeling platform. Proc Natl Acad Sci U S A 119, e2203027119. 10.1073/pnas.2203027119.

Obermann, J., Priglinger, C.S., Merl-Pham, J., Geerlof, A., Priglinger, S., Gotz, M., and Hauck, S.M. (2017). Proteome-wide Identification of Glycosylation-dependent Interactors of Galectin-1 and Galectin-3 on Mesenchymal Retinal Pigment Epithelial (RPE) Cells. Mol Cell Proteomics 16, 1528–1546. 10.1074/mcp.M116.066381.

Pommerenke, C., Rand, U., Uphoff, C.C., Nagel, S., Zaborski, M., Hauer, V., Kaufmann, M., Meyer, C., Denkmann, S.A., Riese, P., et al. (2021). Identification of cell lines CL-14, CL-40 and CAL-51 as suitable models for SARS-CoV-2 infection studies. PLoS One 16, e0255622. 10.1371/journal.pone.0255622.

Poortahmasebi, V., Poorebrahim, M., Najafi, S., Jazayeri, S.M., Alavian, S.M., Arab, S.S., Ghavami, S., Alavian, S.E., Rezaei Moghadam, A., and Amiri, M. (2016). How Hepatitis C Virus Leads to Hepatocellular Carcinoma: A Network-Based Study. Hepat Mon 16, e36005. 10.5812/hepatmon.36005.

Qin, W., Cho, K.F., Cavanagh, P.E., and Ting, A.Y. (2021). Deciphering molecular interactions by proximity labeling. Nat Methods 18, 133–143. 10.1038/s41592-020-01010-5.

Ripa, I., Andreu, S., Lopez-Guerrero, J.A., and Bello-Morales, R. (2021). Membrane Rafts: Portals for Viral Entry. Front Microbiol 12, 631274. 10.3389/fmicb.2021.631274.

Ruiz-Garcia, A., Lopez-Lopez, S., Garcia-Ramirez, J.J., Baladron, V., Ruiz-Hidalgo, M.J., Lopez-Sanz, L., Ballesteros, A., Laborda, J., Monsalve, E.M., and Diaz-Guerra, M.J. (2016). The Tetraspanin TSPAN33 Controls TLR-Triggered Macrophage Activation through Modulation of NOTCH Signaling. J Immunol 197, 3371–3381. 10.4049/jimmunol.1600421.

Rupp, J.C., Locatelli, M., Grieser, A., Ramos, A., Campbell, P.J., Yi, H., Steel, J., Burkhead, J.L., and Bortz, E. (2017). Host Cell Copper Transporters CTR1 and ATP7A are important for Influenza A virus replication. Virol J 14, 11. 10.1186/s12985-016-0671-7.

Ryan, M.J., Johnson, G., Kirk, J., Fuerstenberg, S.M., Zager, R.A., and Torok-Storb, B. (1994). HK-2: an immortalized proximal tubule epithelial cell line from normal adult human kidney. Kidney Int 45, 48–57. 10.1038/ki.1994.6.

Ryu, K.A., Kaszuba, C.M., Bissonnette, N.B., Oslund, R.C., and Fadeyi, O.O. (2021). Interrogating biological systems using visible-light-powered catalysis. Nature Reviews Chemistry 5, 322–337. 10.1038/s41570-021-00265-6.

Scalise, M., Pochini, L., Console, L., Losso, M.A., and Indiveri, C. (2018). The Human SLC1A5 (ASCT2) Amino Acid Transporter: From Function to Structure and Role in Cell Biology. Front Cell Dev Biol 6, 96. 10.3389/fcell.2018.00096.

Shilts, J., Crozier, T.W.M., Greenwood, E.J.D., Lehner, P.J., and Wright, G.J. (2021). No evidence for basigin/CD147 as a direct SARS-CoV-2 spike binding receptor. Sci Rep 11, 413. 10.1038/s41598-020-80464-1.

Srivastava, M., Zhang, Y., Chen, J., Sirohi, D., Miller, A., Zhang, Y., Chen, Z., Lu, H., Xu, J., Kuhn, R.J., and Andy Tao, W. (2020). Chemical proteomics tracks virus entry and uncovers NCAM1 as Zika virus receptor. Nat Commun 11, 3896. 10.1038/s41467-020-17638-y.

Stehle, T., and Dermody, T.S. (2004). Structural similarities in the cellular receptors used by adenovirus and reovirus. Viral Immunol 17, 129–143. 10.1089/0882824041310621.

Suprewicz, L., Swoger, M., Gupta, S., Piktel, E., Byfield, F.J., Iwamoto, D.V., Germann, D., Reszec, J., Marcinczyk, N., Carroll, R.J., et al. (2021). Extracellular vimentin as a target against SARS-CoV-2 host cell invasion. bioRxiv. 10.1101/2021.01.08.425793.

Swaine, T., and Dittmar, M.T. (2015). CDC42 Use in Viral Cell Entry Processes by RNA Viruses. Viruses 7, 6526–6536. 10.3390/v7122955.

Taban, Q., Mumtaz, P.T., Masoodi, K.Z., Haq, E., and Ahmad, S.M. (2022). Scavenger receptors in host defense: from functional aspects to mode of action. Cell Commun Signal 20, 2. 10.1186/s12964-021-00812-0.

Teo, G., Liu, G., Zhang, J., Nesvizhskii, A.I., Gingras, A.C., and Choi, H. (2014). SAINTexpress: improvements and additional features in Significance Analysis of INTeractome software. J Proteomics 100, 37–43. 10.1016/j.jprot.2013.10.023.

Uhlen, M., Fagerberg, L., Hallstrom, B.M., Lindskog, C., Oksvold, P., Mardinoglu, A., Sivertsson, A., Kampf, C., Sjostedt, E., Asplund, A., et al. (2015). Proteomics. Tissue-based map of the human proteome. Science 347, 1260419. 10.1126/science.1260419.

V’Kovski, P., Gerber, M., Kelly, J., Pfaender, S., Ebert, N., Braga Lagache, S., Simillion, C., Portmann, J., Stalder, H., Gaschen, V., et al. (2019). Determination of host proteins composing the microenvironment of coronavirus replicase complexes by proximity-labeling. Elife 8. 10.7554/eLife.42037.

Vanderbeck, A., and Maillard, I. (2021). Notch signaling at the crossroads of innate and adaptive immunity. J Leukoc Biol 109, 535–548. 10.1002/JLB.1RI0520-138R.

Verschueren, E., Von Dollen, J., Cimermancic, P., Gulbahce, N., Sali, A., and Krogan, N.J. (2015). Scoring Large-Scale Affinity Purification Mass Spectrometry Datasets with MiST. Curr Protoc Bioinformatics 49, 8 19 11–18 19 16. 10.1002/0471250953.bi0819s49.

Walls, A.C., Park, Y.J., Tortorici, M.A., Wall, A., McGuire, A.T., and Veesler, D. (2020). Structure, Function, and Antigenicity of the SARS-CoV-2 Spike Glycoprotein. Cell 183, 1735. 10.1016/j.cell.2020.11.032.

Wang, H., Li, Z.Y., Liu, Y., Persson, J., Beyer, I., Moller, T., Koyuncu, D., Drescher, M.R., Strauss, R., Zhang, X.B., et al. (2011). Desmoglein 2 is a receptor for adenovirus serotypes 3, 7, 11 and 14. Nat Med 17, 96–104. 10.1038/nm.2270.

Xu, Z., and Jin, B. (2010). A novel interface consisting of homologous immunoglobulin superfamily members with multiple functions. Cell Mol Immunol 7, 11–19. 10.1038/cmi.2009.108.

Yeung, M.L., Teng, J.L.L., Jia, L., Zhang, C., Huang, C., Cai, J.P., Zhou, R., Chan, K.H., Zhao, H., Zhu, L., et al. (2021). Soluble ACE2-mediated cell entry of SARS-CoV-2 via interaction with proteins related to the renin-angiotensin system. Cell 184, 2212–2228 e2212. 10.1016/j.cell.2021.02.053.

Zhang, F., Li, W., Feng, J., Ramos da Silva, S., Ju, E., Zhang, H., Chang, Y., Moore, P.S., Guo, H., and Gao, S.J. (2021). SARS-CoV-2 pseudovirus infectivity and expression of viral entry-related factors ACE2, TMPRSS2, Kim-1, and NRP-1 in human cells from the respiratory, urinary, digestive, reproductive, and immune systems. J Med Virol 93, 6671–6685. 10.1002/jmv.27244.

Zhang, H., Li, Y., Wang, H.B., Zhang, A., Chen, M.L., Fang, Z.X., Dong, X.D., Li, S.B., Du, Y., Xiong, D., et al. (2018). Ephrin receptor A2 is an epithelial cell receptor for Epstein-Barr virus entry. Nat Microbiol 3, 1–8. 10.1038/s41564-017-0080-8.

Zhang, Y., Shang, L., Zhang, J., Liu, Y., Jin, C., Zhao, Y., Lei, X., Wang, W., Xiao, X., Zhang, X., et al. (2022). An antibody-based proximity labeling map reveals mechanisms of SARS-CoV-2 inhibition of antiviral immunity. Cell Chem Biol 29, 5–18 e16. 10.1016/j.chembiol.2021.10.008.

Zheng, K.I., Feng, G., Liu, W.Y., Targher, G., Byrne, C.D., and Zheng, M.H. (2021). Extrapulmonary complications of COVID-19: A multisystem disease? J Med Virol 93, 323–335. 10.1002/jmv.26294.

Zhou, P., Yang, X.L., Wang, X.G., Hu, B., Zhang, L., Zhang, W., Si, H.R., Zhu, Y., Li, B., Huang, C.L., et al. (2020). A pneumonia outbreak associated with a new coronavirus of probable bat origin. Nature 579, 270–273. 10.1038/s41586-020-2012-7.

