## Supplementary material for "High resolution photocatalytic mapping of SARS-CoV-2 Spike protein-host cell membrane interactions": ViraMap Supplementary Information

**This PDF file includes:**

Figures S1 to S4
Legends for Figures S1 to S4

**Other supplementary materials for this manuscript include the following:**

Table S1 to S5 (titles are included in the main text manuscript file)

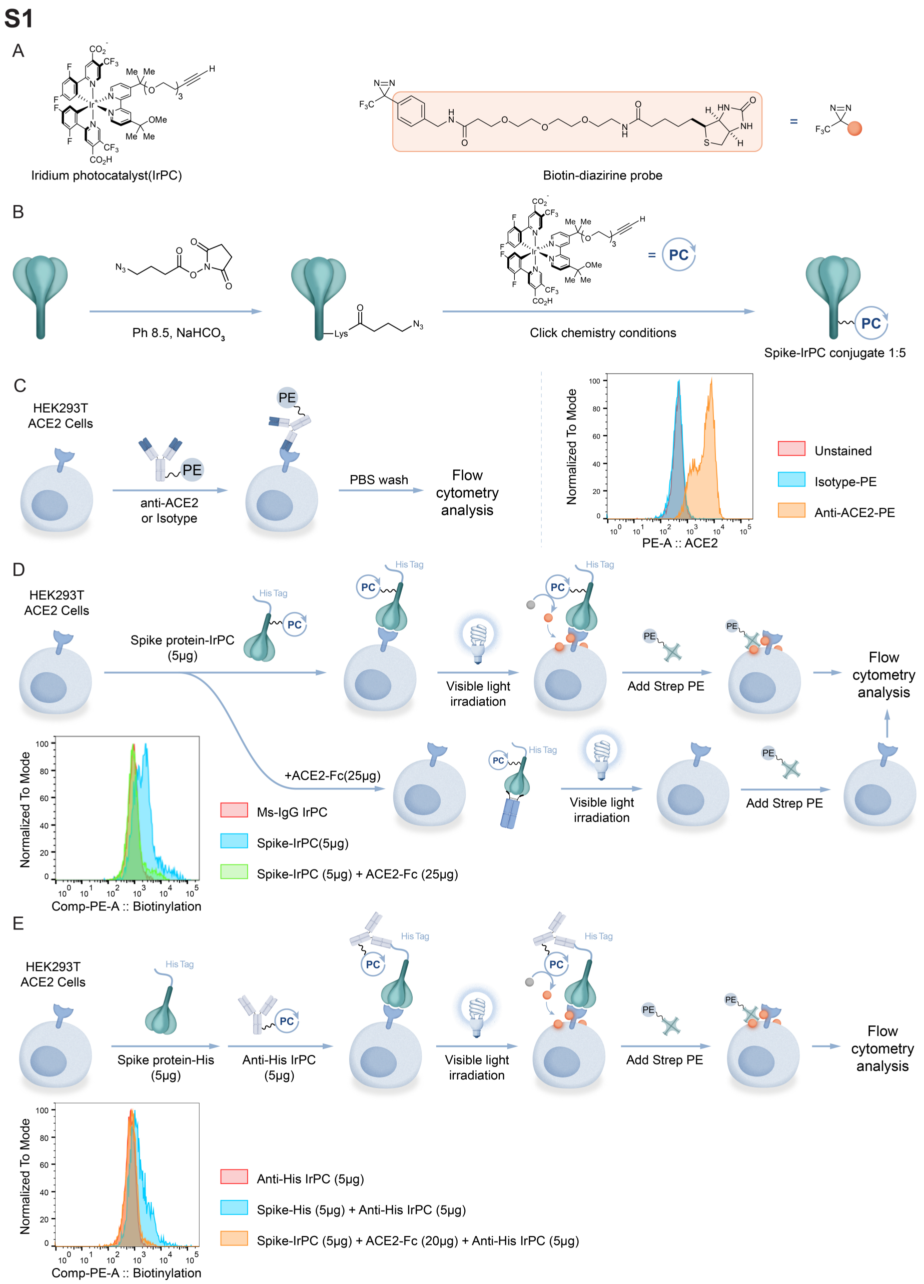


**Figure S1.** A. Structure of the Iridium photocatalyst (IrPC) and diazirine biotin probe used in this work. B. Preparation of Spike-IrPC trimer conjugate. C. Surface detection of ACE2 on HEK293T+ACE2 overexpression cells. D. Free ACE2-Fc binding by Spike-IrPC in HEK293T+ACE2 cells confirmed ACE2 specific binding function of the Spike-IrPC conjugate. E. Surface biotinylation of HEK293T+ACE2 cells using IrPc on anti-His antibody probing the His-tagged Spike trimer.

**S2**

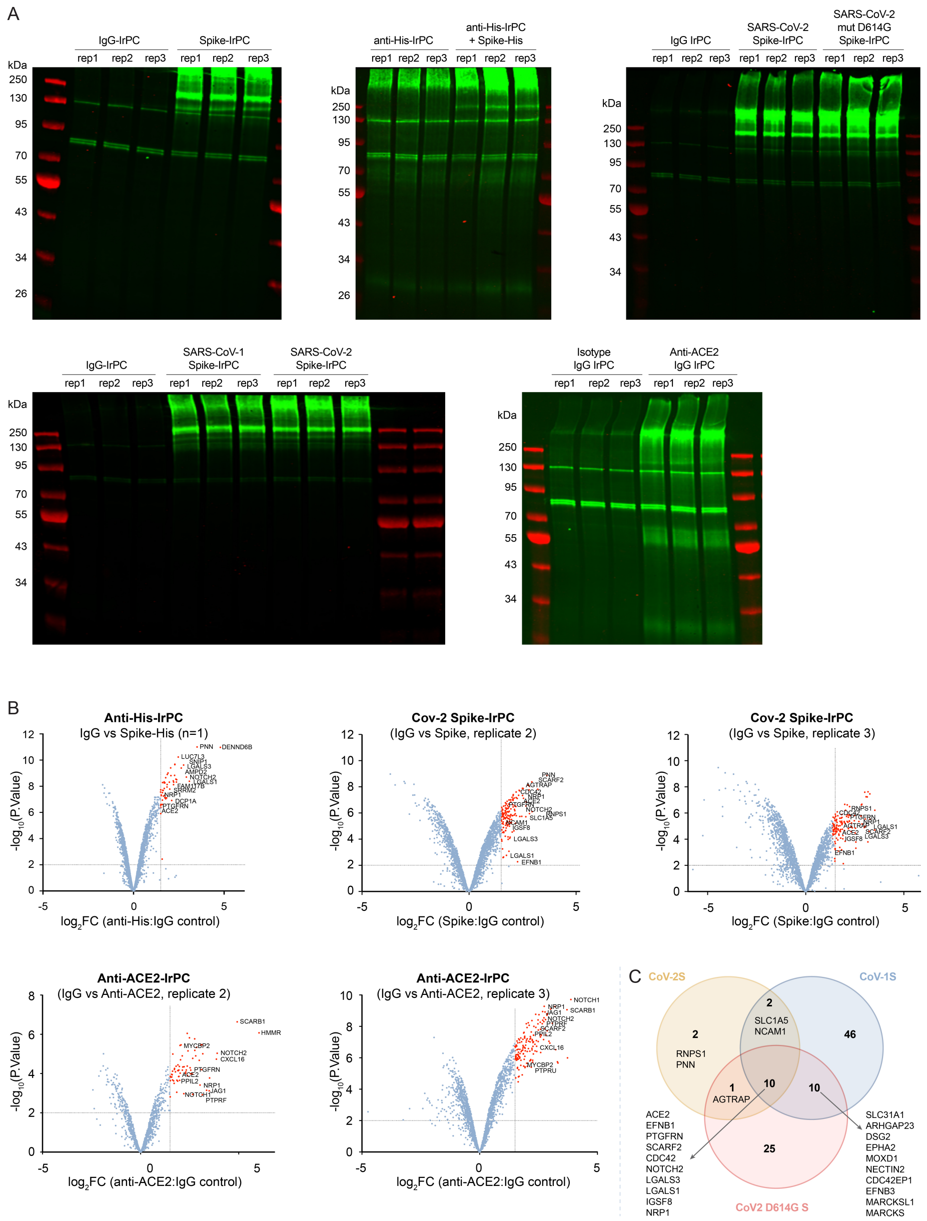
A
IgG-IrPC
Spike-IrPC
rep1 rep2 rep3 rep1 rep2 rep3
kDa
250
130
95
70
55
43
34
26
anti-His-IrPC
anti-His-IrPC + Spike-His
IgG IrPC
SARS-CoV-2 Spike-IrPC
SARS-CoV-2 mut D614G Spike-IrPC
SARS-CoV-1 Spike-IrPC
Isotype IgG IrPC
Anti-ACE2 IgG IrPC
B
Anti-His-IrPC
IgG vs Spike-His (n=1)
log2FC (anti-His:IgG control)
Cov-2 Spike-IrPC
(IgG vs Spike, replicate 2)
(IgG vs Spike, replicate 3)
log2FC (Spike:IgG control)
-log10(P.Value)
Anti-ACE2-IrPC
(IgG vs Anti-ACE2, replicate 2)
(IgG vs Anti-ACE2, replicate 3)
log2FC (anti-ACE2:IgG control)
C
CoV-2S
CoV-1S
CoV2 D614G S
2
SLC1A5
NCAM1
2
RNPS1
PNN
46
1
AGTRAP
10
10
25
ACE2
EFNB1
PTGFRN
SCARF2
CDC42
NOTCH2
LGALS3
LGALS1
IGSF8
NRP1
SLC31A1
ARHGAP23
DSG2
EPHA2
MOXD1
NECTIN2
CDC42EP1
EFNB3
MARCKSL1
MARCKS

## S2-2

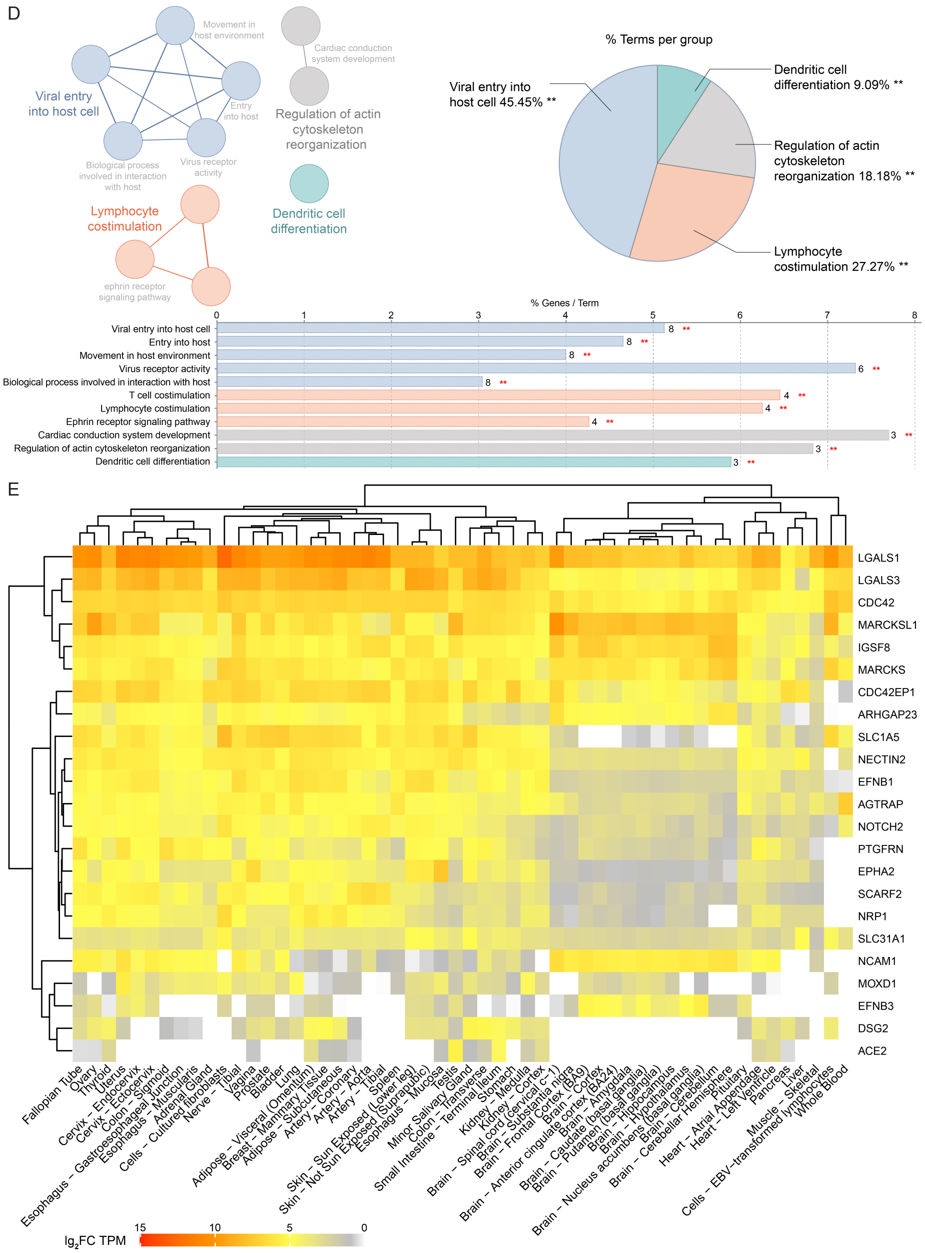


**Figure S2.** A. Detection of the biotinylated fraction of HEK293T+ACE2 cells after targeted labeling using IrPC-labeled target proteins by Western blotting. B. Volcano plots showing enriched host factors from anti-His targeted labeling and replicate experiments of SARS-CoV-2 Spike and ACE2 targeted labeling. C. Overlap of enriched proteins obtained from ViraMapping with the different Spike-IrPC variants. D. Gene-ontology enrichment analysis of the 23 highconfidence (HC) enriched proteins using Cytoscape 3.9.1 plugin ClueGO classified the interactors into 11 terms/pathways and 4 functional groups (Ontology used: GO_BiologicalProcess-EBI-UniProt-GOA-ACAP-ARAP_13.05.2021_00h00) (genes associated with each pathway are indicated in Table S2). E. Heatmap of tissue expression profiles of the targeted labeling-output list of 23 HC enriched proteins.

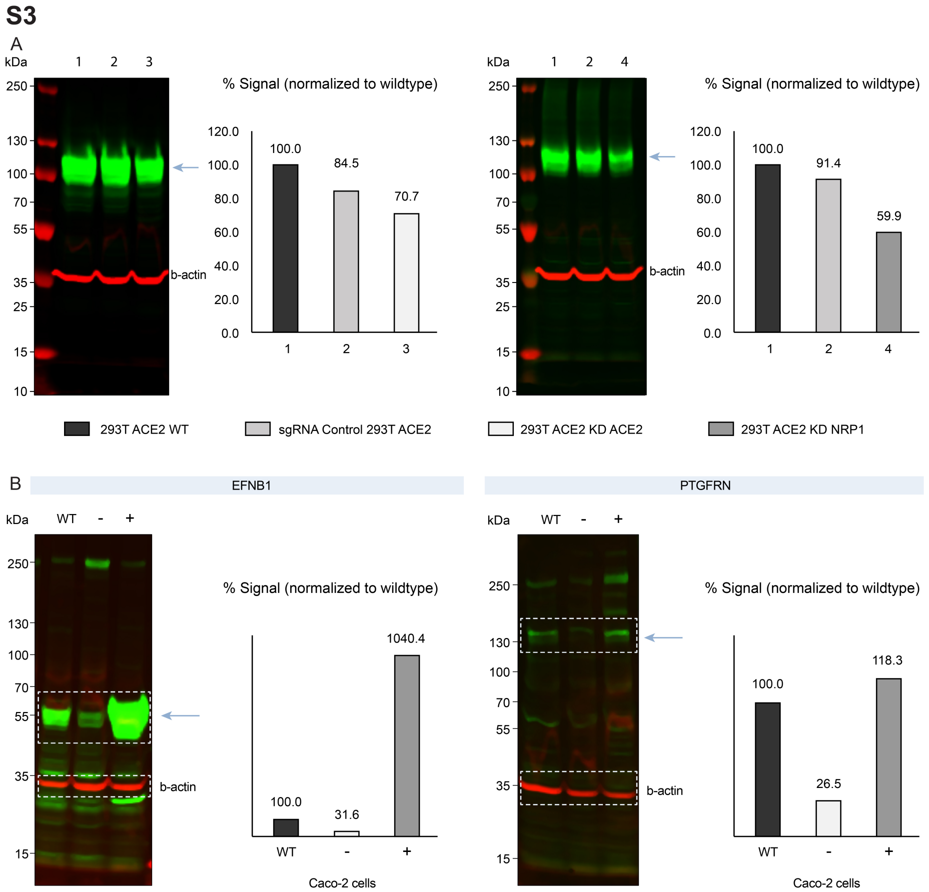


**Figure S3**. A. CRISPR-Cas9 mediated knockdown efficiency of ACE2 (~30%) and NRP1 (~40%) in HEK293T+ACE2 cells. B. Confirmation of knockdown and over-expression of EFNB1 and PTGFRN in Caco-2 cells by immunoblotting and scanning under Li-Cor CLx infrared scanner. Each band size of target proteins matched with the claimed band size on the manufacturer's website. Multiple bands were observed at various molecular weights for these proteins in Caco-2 cells, possibly due to a combination of factors such as different isoforms of the same protein and/or cross-reactivity of similar epitopes on other proteins by the primary antibodies for this cell type. A slightly higher molecular weight of EFNB1 on the immunoblots was observed indicating possible post-translational modifications.

S4

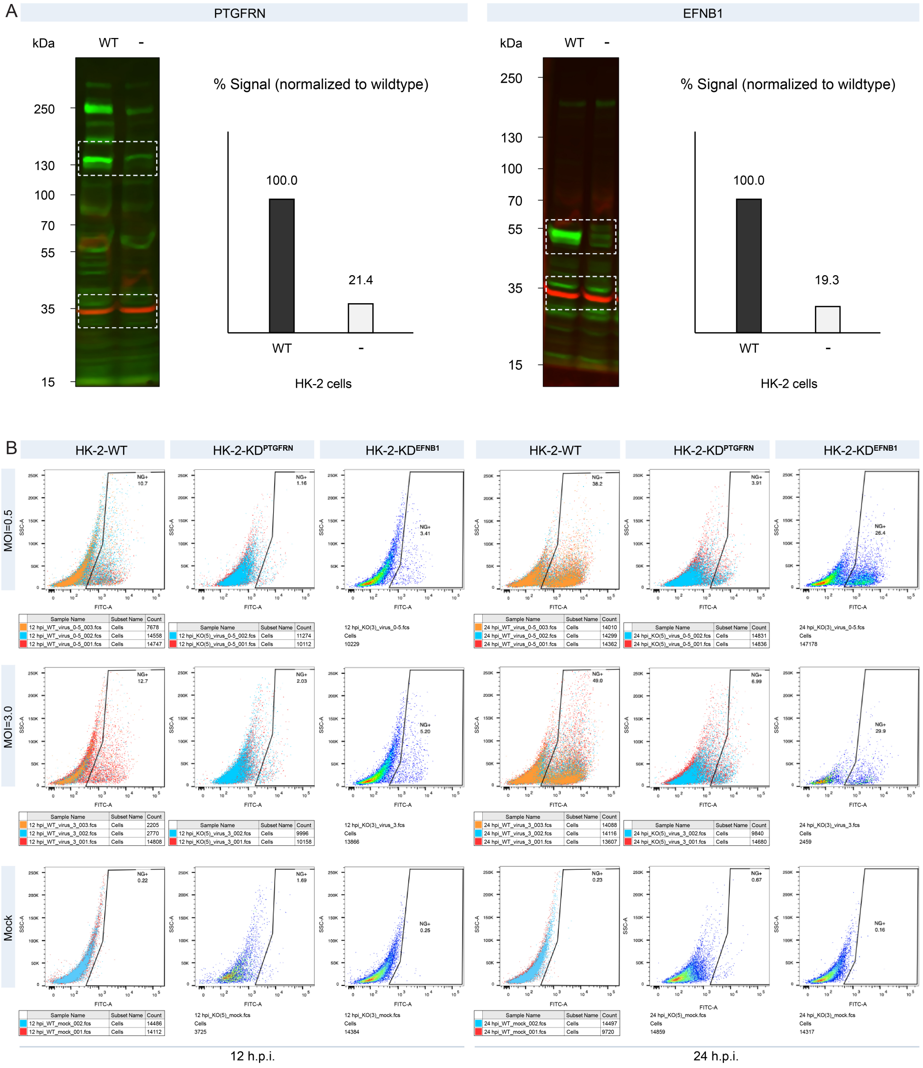
A
PTGFRN
kDa
WT
-
250
130
100
70
55
35
15
% Signal (normalized to wildtype)
100.0
21.4
WT
-
HK-2 cells
EFNB1
kDa
WT
-
250
130
100
70
55
35
15
% Signal (normalized to wildtype)
100.0
19.3
WT
-
HK-2 cells
B
HK-2-WT
HK-2-KDPTGFRN
HK-2-KDEFNB1
HK-2-WT
HK-2-KDPTGFRN
HK-2-KDEFNB1
MOI=0.5
MOI=3.0
Mock
FITC-A
SSC-A
12 h.p.i.
24 h.p.i.

70

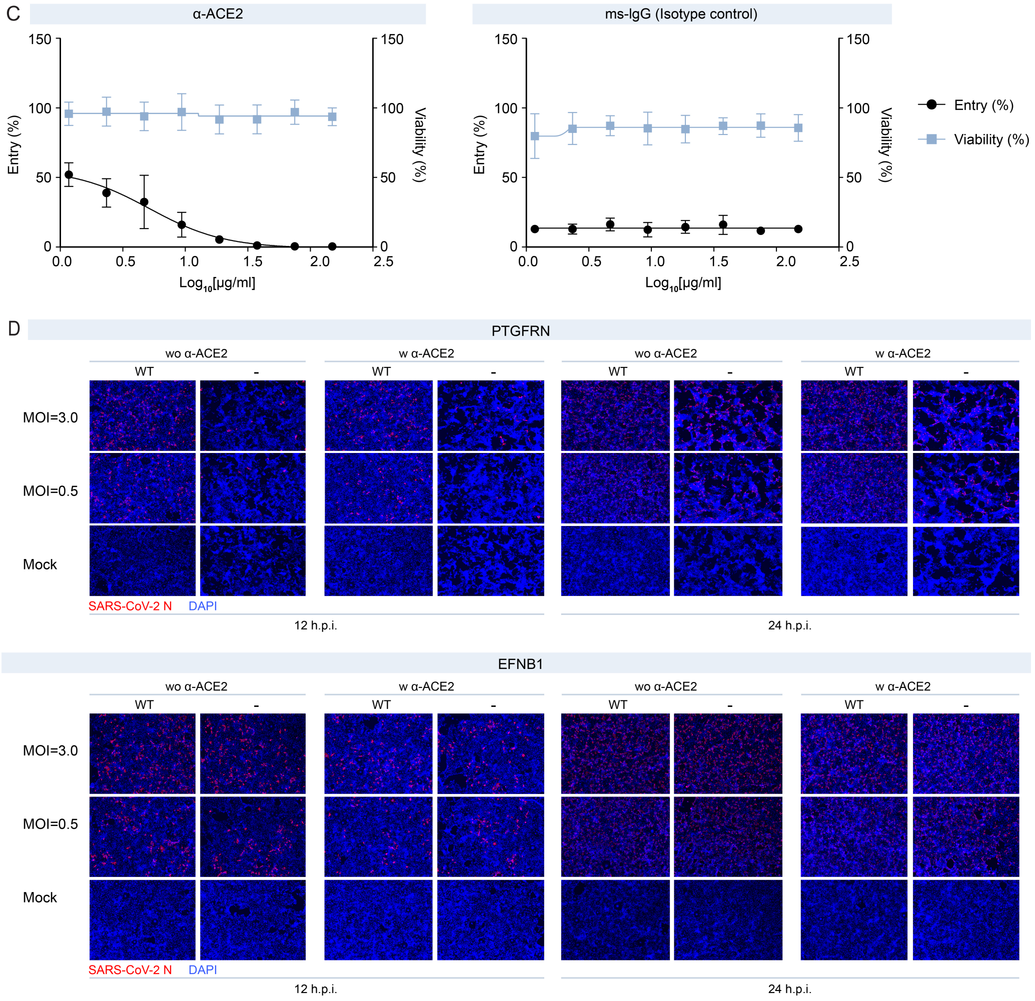


**Figure S4.** A. Immunoblots for PTGFRN and EFNB1 expression knockdown in HK-2 cells. B. Flow cytometry analysis showing percentage of infected cell population in wildtype (WT), PTGFRN and EFNB1 knockdown HK2 cells upon SARS-CoV-2 infection at different multiplicity of infection (0.5 and 3.0) and time points (12 h.p.i. and 24 h.p.i.) of infection. C. Determination of $[ACE2_{50}]$ for HK-2 wildtype cells infected by CoV-2-S pseudotyped virus. D. Representative confocal images of SARS CoV-2 infected wildtype (WT), PTGFRN and EFNB1 knockdown HK-2 cells at different multiplicity of infection (0.5 and 3.0) and time points of infection (12 h.p.i. and 24 h.p.i.) in presence and absence of ACE2-blockade.
